# A non-mosaic humanized mouse model of Down syndrome, trisomy of a nearly complete long arm of human chromosome 21 in mouse chromosome background

**DOI:** 10.1101/862433

**Authors:** Yasuhiro Kazuki, Feng J. Gao, Yicong Li, Anna J. Moyer, Benjamin Devenney, Kei Hiramatsu, Sachiko Miyagawa-Tomita, Satoshi Abe, Kanako Kazuki, Naoyo Kajitani, Narumi Uno, Shoko Takehara, Masato Takiguchi, Miho Yamakawa, Atsushi Hasegawa, Ritsuko Shimizu, Satoko Matsukura, Naohiro Noda, Narumi Ogonuki, Kimiko Inoue, Shogo Matoba, Atsuo Ogura, Liliana D. Florea, Alena Savonenko, Meifang Xiao, Dan Wu, Denise A.S. Batista, Junhua Yang, Zhaozhu Qiu, Nandini Singh, Joan T. Richstemeier, Takashi Takeuchi, Mitsuo Oshimura, Roger H. Reeves

## Abstract

Down syndrome (DS) is a complex human condition, and animal models trisomic for human chromosome 21 (HSA21) genes or orthologs provide insights into better understanding and treating DS. However, HSA21 orthologs are distributed into three mouse chromosomes, preventing us from generating mouse models trisomy of a complete set of HSA21 orthologs. The only existing humanized mouse DS model, Tc1, carries a HSA21 with over 20% of protein coding genes (PCGs) disrupted. More importantly, due to the human centromere, Tc1 is mosaic (a mix of euploid and trisomic cells), which makes every mouse unique and compromises interpretation of results. Here, we used mouse artificial chromosome (MAC) technology to “clone” the 34 MB long arm of HSA21 (HSA21q). Through multiple steps of microcell-mediated chromosome transfer we created a new humanized DS mouse model, Tc(HSA21q;MAC)1Yakaz (“TcMAC21”). Constitutive EGFP expression from the transchromosome and fluorescent in situ hybridization validate that TcMAC21, containing a hybrid chromosome of HSA21q and mouse centromere, is not mosaic. Whole genome sequencing shows that TcMAC21 contains a nearly complete copy of HSA21q with 93% of intact PCGs, while RNA-seq and additional mRNA/protein expression analyses confirm that PCGs are transcribed and regulated. A battery of tests show that TcMAC21 recapitulates many DS phenotypes including morphological anomalies in heart, craniofacial skeleton and brain, pathologies at molecular and cellular level, and impairments in learning, memory and synaptic plasticity. TcMAC21 is the most complete mouse model of DS extant and has potential for supporting a wide range of basic and preclinical research.

**Significance Statement:** In the last 25 years, mouse models of trisomy 21 have supported research into Down syndrome, from defining the basis for developmental effects up to support for clinical trials. However, existing models have significant shortfalls, especially for preclinical studies. These deficiencies include incomplete or inappropriate representation of trisomic genes, absence of an extra chromosome, and mosaicism.

Using cutting edge technologies we produced a mouse artificial chromosome containing the entire 34Mb long arm of human chromosome 21 and, with assisted reproductive technologies, established it in the germ line of mice. This trisomic mouse manifests developmental and functional features of Down syndrome, including hippocampal-based learning and memory deficits. This is the most complete model of Down syndrome produced to date.

## INTRODUCTION

Human aneuploidy, a gain or loss of chromosomes, is associated with both birth defects and cancer(1). Because of mitotic abnormalities and hundreds of genes having dosage imbalance, most of aneuploidies lead to miscarriage. Down syndrome (DS), also known as trisomy 21, is the most common aneuploidy in live births, occurring in about 1 in every 800 babies (2). Besides common facial and other physical features, People with DS have intellectual disabilities, congenital heart disease, higher risk of leukemia, early onset dementia and so on (3). The impact of trisomic mouse models on DS research cannot be overestimated. Davisson’s bold experiment to create translocations of mouse chromosomes generated the first viable DS model, Ts65Dn (4, 5). The demonstration that features directly comparable to those in DS occurred in mice with trisomy for a number of genes orthologous to human chromosome 21 (HSA21) completely changed the paradigm for research in this area (6). Ts65Dn has been the most broadly used model for DS research for more than two decades, up to and including crucial support for clinical trials (7). However, advances in genomics have defined HSA21 and its mouse orthologs much more precisely, and a better genetic representation is necessary to support current research efforts (8–11).

There are a number of technical arguments for a new DS model to meet these challenges, but perhaps the most compelling global issue relates to pre-clinical drug testing. Potential treatments in Ts65Dn mice have yielded an embarrassment of riches with at least a dozen potential single gene targets whose ablation alleviates cognitive deficits and more than a dozen pharmaceuticals or nutriceuticals reporting promising effects (12). For example, the recent clinical trial by Roche of the GABA-α5 antagonist RG1662 gave serious weight to experiments in Ts65Dnand other trisomic mice showing that normalization of the balance of inhibitory and excitatory inputs to hippocampus could improve several hippocampal-based behavioral paradigms as well as long term potentiation (LTP), all of which are impaired in Ts65Dn and other DS models (7, 13–15). Improvements with RG1662 are dramatic in mice, but the human trial was terminated in Phase IIa due to lack of efficacy in a long term treatment paradigm, possibly due to accommodation (https://clinicaltrials.gov/ct2/show/NCT02024789). The degree to which the failure to translate a preclinical result to human clinical trials relates to the model itself is unknown. However, it seems reasonable to expect that a better genetic representation of trisomy 21 will produce more relevant results.

Tc1 is the only existing DS mouse model carrying a HSA21, but more than 50 of protein coding genes (PCGs) in HSA21q were disrupted by rearrangement, deletion or duplication (16, 17). A further limitation of Tc1 is that all mice are mosaic for HSA21, that is, the human chromosome is randomly lost from cells during mitosis. Mosaicism makes each Tc1 mouse unique from every other mouse, because the developmental trajectories dependent on cell-cell interactions are different in every individual (18, 19). Mosaicism in Tc1 is not an isolated case, as the retention rates of other human chromosomes and human artificial chromosomes (HACs) vary widely in different mouse tissues (20–22). The centromere is a multi-function regulator of genome stability and linked to the cause of chromosomal instability found in the interspecific hybridization (23, 24).

To overcome the problem of mosaicism, we developed mouse artificial chromosome vectors (MACs), capable of carrying Mb-sized genomic segments. MACs are freely segregating and stably maintained in mice (22, 25). Here we report the transfer of a nearly complete copy of HSA21q into the mouse chromosome background using a MAC-mediated genomic transfer (MMGT) system. The new humanized mouse model of DS, TcMAC21, shows no mosaicism. Whole genome sequencing, RNA-Seq, protein expression, and phenotypic analyses confirm that TcMAC21 has the most complete representation of HSA21 genes among all DS mouse models, and that a number of phenotypes mirroring those common in individuals with DS occur in TcMAC21.

## RESULTS

### Construction and whole genome sequencing (WGS) of TcMAC21 mice

We previously constructed hybrid A9 cells containing a copy of HSA21 (26), which was then moved into DT40 cells by microcell-mediated chromosome transfer (MMCT) (27). A loxP site was introduced into the HSA21 NC_000021.9 locus (13021348-13028858) to create HSA21-loxP. In this study, the HSA21-loxP chromosome was transferred by MMCT into Chinese hamster ovary (CHO) cells containing the MAC1 vector, which includes EGFP, a neomycin resistance gene, and loxP (Fig. 1A **and Fig. S1, A and B)**. To induce reciprocal translocation between MAC1 and HSA21-loxP, a Cre-expressing vector was transfected into the hybrid cells, reconstituting *HPRT* to allow selection of recombinant clones in HAT medium. Two lines were confirmed by FISH to contain the HSA21q-MAC **(Fig. S1C)**, and the HSA21q-MAC from each line was introduced into TT2F female mouse ES cell lines using MMCT. FISH confirmed that two of the five lines contained HSA21q-MAC as a freely segregating chromosome **(Fig. S1D)**. The two FISH-positive TT2F clones with karyotypes of 40, XO, +HSA21q-MAC were used to produce chimeras that had various degrees of coat-color chimerism (Fig. 1B). Six GFP-positive female chimeras were crossed with ICR males to establish the TcMAC21 mouse strain and a line stably segregating the HSA21q-MAC was recovered. Both FISH and G-banding identified the supernumerary artificial chromosome in TcMAC21, and G-banding of HSA21q-MAC1 showed HSA21q-like pattern (Fig. 1, C and D). TcMAC21 was crossed for eight generations onto BDF1 strain background and transferred to the Riken Animal Resource (BRC No. RBRC05796 and STOCK Tc (HSA21q-MAC1)). An inbred subline (B6. TcMAC21) has been initiated by crossing onto C57BL/6J (B6) and was used for some of the phenotypic characterizations presented here as indicated. WGS revealed that TcMAC21 had a nearly complete HSA21q, covering all PCGs from POTED to PRMT2 with four deletions in between (Fig. 1E**, and Data file S1)**. While physical deletions encompass ∼29% of HSA21q, they occur substantially in gene-poor regions. Consequently, only 14 out of 213 HSA21q PCGs were deleted(Fig. 1F).

**Fig. 1.**
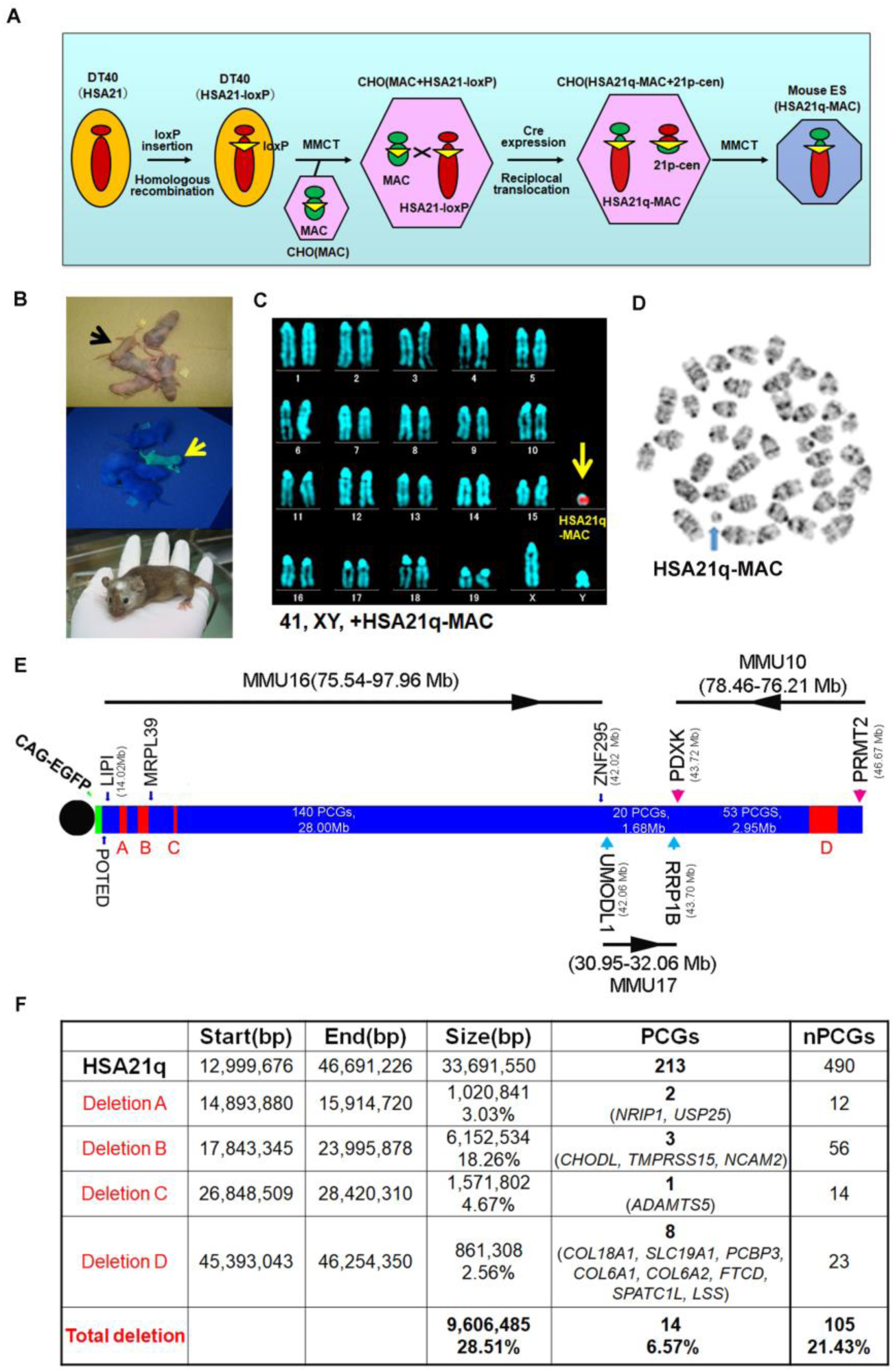
Construction of TcMAC21 mice (HSA21q-MAC). (**A**) Schematic diagram of HSA21q-MAC construction. (**B**) Chimeric mice were obtained via injection of mouse ES cells carrying the HSA21q-MAC. The arrow indicates a GFP-positive, TcMAC21 trisomic mouse. (**C**) FISH and (**D**) G-banding based karyotype of TcMAC21 containing the HSA21q-MAC. (**E**) WGS showing the positions of 4 deletions in HSA21q (A, B, C, and D). These are shown normalized to gene numbers, not physical length. The regions of homology with mouse chromosomes 16, 17 and 10 are indicated. (**F**) Genome positions of deletions and numbers of affected PCGs and non-PCGs on the HSA21q-MAC (based on GRCh38.p12, BioMart-Ensembl, May 2019).

### HSA21 genes are expressed, producing dosage imbalance in TcMAC21 mice

We used RNA-Seq to determine whether HSA21q genes are expressed by comparing forebrain mRNA levels between TcMAC21 and euploid (Eu) at postnatal day 1 (P1). As expected, neither 49 HSA21q keratin associated protein genes (KRTAPs) nor their mouse orthologs were expressed in TcMAC21 or Eu brain **(Data file S2)**. Excluding KRTAPs, 160 PCGs on HSA21q have mouse orthologs. We compared TcMAC21 with Eu gene expression from the 117 mouse orthologs with FPKM ≥1 in Eu (Fig. 2A). Expression of mouse orthologs was not substantially affected by the presence of HSA21q in TcMAC21, as 106 out of 117 mouse orthologs in TcMAC21 were expressed at 80-120% of Eu levels. Two mouse orthologs, *Erg* and *Prdm15*, had reduced expression (<0.8-fold) and nine genes showed increased mouse ortholog expression (>1.2-fold) in TcMAC21.

**Fig. 2.**
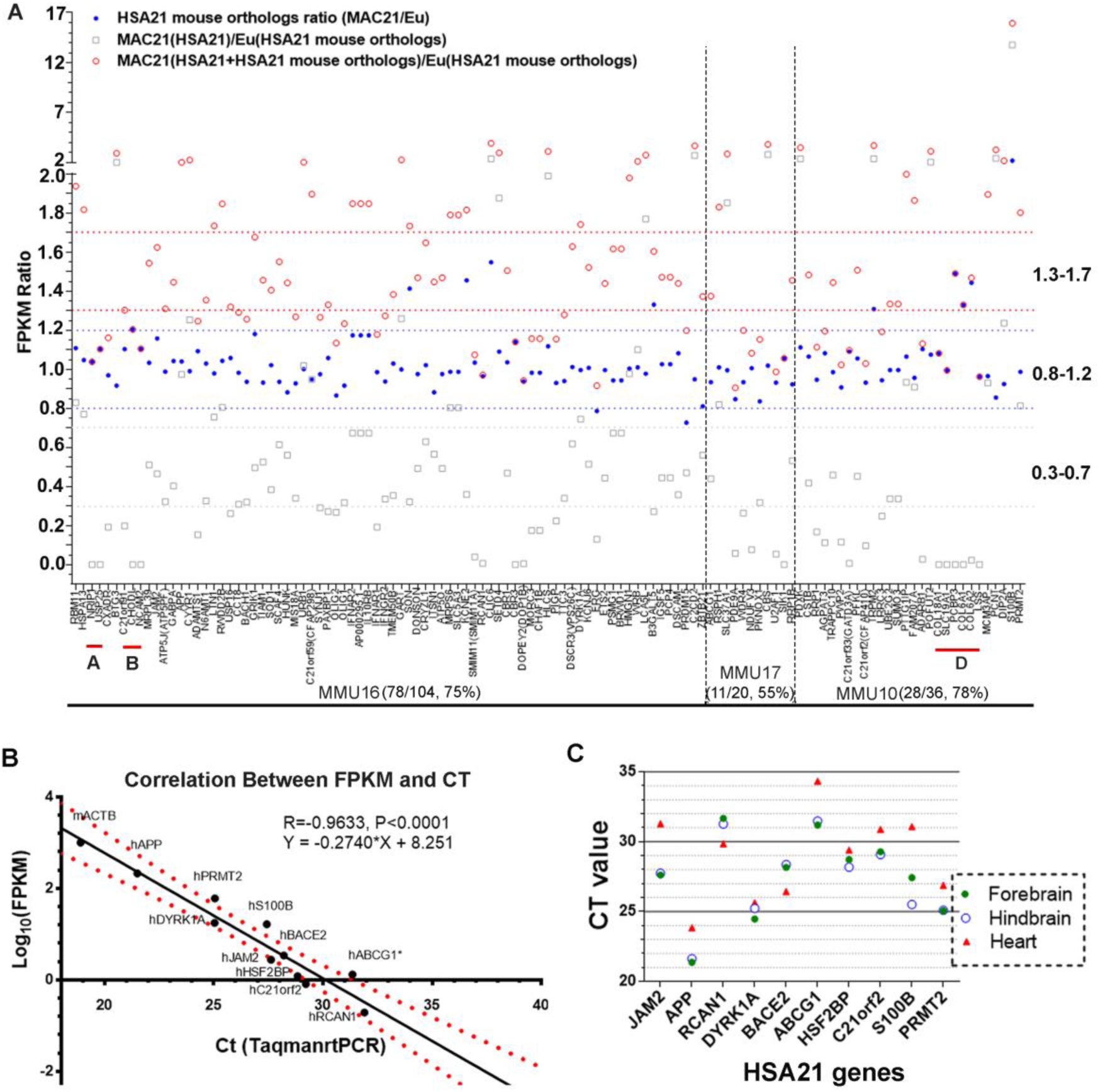
HSA21q PCG expression pattern in P1 TcMAC21. (**A**) RNA-Seq analysis of HSA21q and its mouse orthologs in TcMAC21 and Eu in P1 forebrain identified 117 HSA21 genes with mouse orthologs whose FPKM is ≥1 in Eu. Three expression values are shown: 1. the ratio of HSA21q transcript in TcMAC21 relative to its ortholog in Eu mice (gray open squares); 2. the ratio of the mouse ortholog in TcMAC21 relative to that of Eu (blue circles); and 3. the ratio of total expression (FPKM of HSA21q + its mouse ortholog) in TcMAC21 relative to Eu (red circles). The positions of deleted regions are indicated in red. (**B**) Correlation between relative quantification by Taqman assay and by RNA-Seq for 10 HSA21q genes and mouse actin (mACTB) in TcMAC21. (**C**) Taqman assay comparing expression of 10 HSA21 genes between forebrain, hindbrain, and heart using the same amount of total RNA. (n = 2 TcMAC21 and 3 Eu littermates at P1).

The mRNA expression levels of HSA21 PCGs were on average about half that of their mouse orthologs, but this HSA21/HSA21 mouse ortholog expression ratio varies considerably across genes (Fig. 2A). We analyzed the ratio of total expression (sum of FPKM for the HSA21 gene + its mouse ortholog) from TcMAC21 to the related mouse ortholog expression level in Eu. As expected, there was considerable variation around a 1.5-fold average increase in expression; 40 genes fell in the low overexpression range (<1.3-fold increased over Eu), 39 genes were in the “expected” range (1.3-1.7-fold), and 38 genes were more highly overexpressed (>1.7-fold). To validate RNA-Seq quantification, 10 human transcript-specific Taqman probes were used to assess forebrain mRNA, and relative expression quantified by Taqman was significantly correlated with that determined by RNA-Seq (R=0.96) (Fig. 2B). Using Taqman assays on total RNA isolated from different tissues of TcMAC21 mice at P1, we found that HSA21 PCG expression levels in forebrain and hindbrain were very similar and frequently different from levels of the same transcript in heart, consistent with tissue-specific regulation of HSA21 genes (Fig. 2C). All results indicate that HSA21q is actively transcribed and regulated to produce dosage imbalance in TcMAC21.

### TcMAC21 mice are not mosaic for trisomy

The peri-centromeric GFP on the HSA21q-MAC allows rapid identification of TcMAC21 by illuminating with UV light (Nightsea). To exclude the possibility that GFP-positive mice could carry a distally deleted chromosome due to translocation or chromosome breakage, we validated the model initially by human specific Taqman assays for ten existing TcMAC21-HSA21q genes plus two HSA21q genes that are absent from TcMAC21 as negative controls. Two TcMAC21 and three Eu littermates showed expected expression patterns indicating that the entire chromosome was retained in TcMAC21 **(Table S1)**. Twelve additional GFP positive (trisomic) and negative (Eu) pairs were analyzed by Taqman for HSA21q proximal and distal markers (*APP* and *PRMT2*, respectively, **Fig. S2A**). Including the TcMAC21 mouse used for WGS, we saw 100% concordance of GFP with HSA21q gene expression across the chromosome and zero false positives among fifteen pairs. Thus, GFP appears to be a reliable marker for genotyping TcMAC21.

At a gross level, we saw no evidence of mosaicism in skin, which would appear as patches of non-fluorescent cells when examined for GFP fluorescence (Fig. 1B **and Fig. S2B)**. Similarly, organs from TcMAC21 mice appeared to be uniformly labeled with GFP (Fig. 3A), and human-specific RT-PCR on RNA from nine organs showed the expected pattern of expression for the eight HSA21 genes tested (Fig. 3B). Several tissues were dissociated and analyzed using FISH, whereby the HSA21q-MAC was detected in ≥96% of TcMAC21 cells (Fig. 3, C and D). Similarly, flow cytometry analysis of lymphocytes showed that over 92% of cells from TcMAC21 were GFP positive (Fig. 3E).

**Fig. 3.**
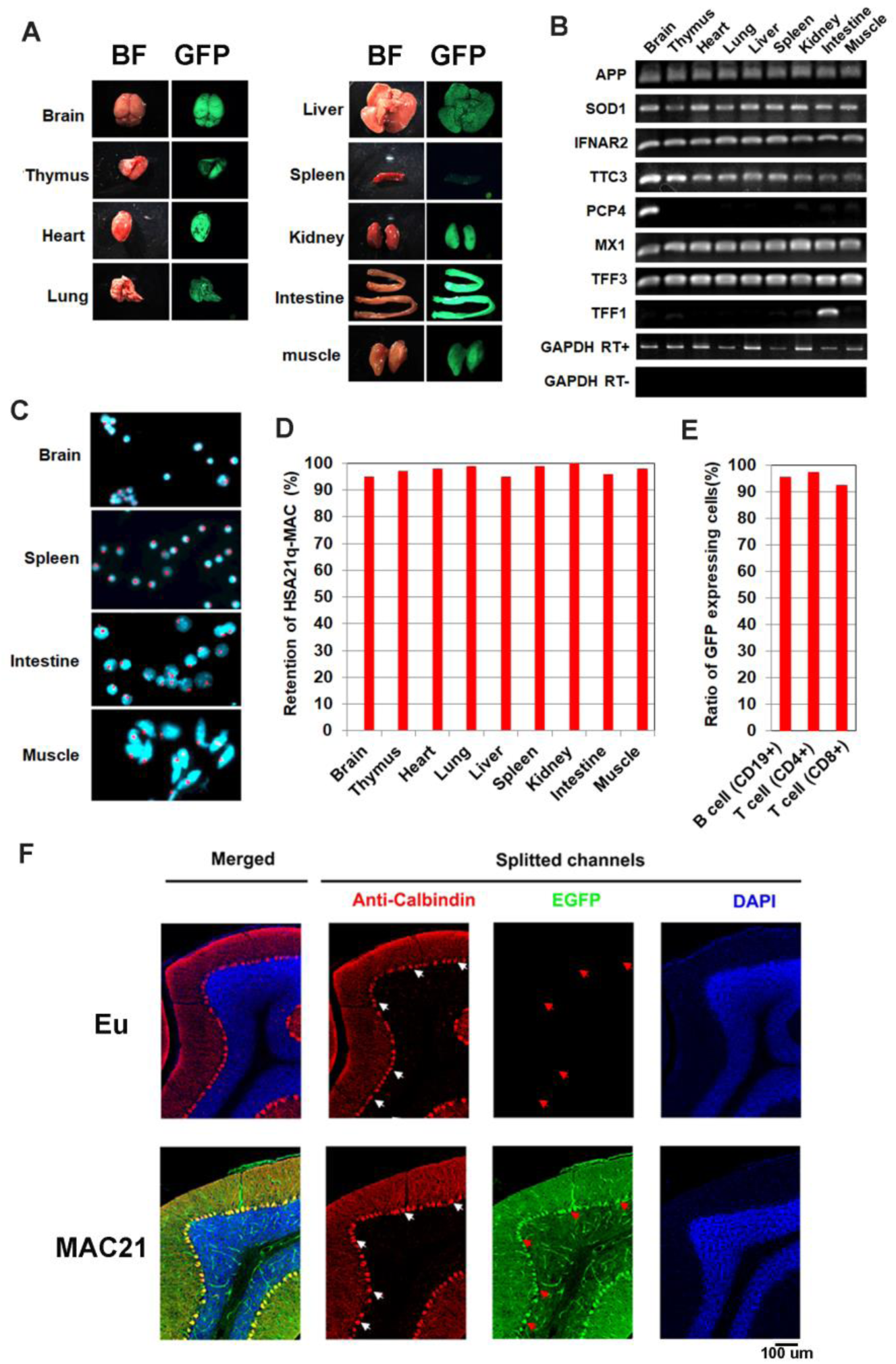
Mosaicism analysis in TcMAC21. (**A**) Nine organs showed uniform GFP expression when illuminated with UV light. (**B**) RT-PCR analyses for HSA21 gene expression in different TcMAC21 tissues demonstrated expected patterns of expression. (**C**) Representative FISH image of cells from TcMAC21 tissues (brain, spleen, intestine and muscle). Red signals indicate the HSA21q-MACs. (**D**) Percentage of cells showing retention of the HSA21q-MAC in various TcMAC21 tissues analyzed by FISH. (**E**) Retention rate of the HSA21q-MAC in three lymphocyte populations analyzed by FCM. (**F**) Mosaicism analysis of HSA21q-MAC in Purkinje cells (PC) of TcMAC21 by immunostaining. White arrows indicate cell bodies of randomly selected PC in RFP-channel, and red arrows indicate corresponding locations in GFP-channel.

Immunostaining of parasagittal brain sections with calbindin showed all Purkinje cells (PCs) in TcMAC21 were GFP positive while those in Eu were GFP negative (Fig. 3F).

Together these findings indicate that, if it occurs at all, mosaicism is rare in TcMAC21. These stable profiles are consistent with those observed previously in mice carrying the empty MAC1 vector (22, 28).

### TcMAC21 has abnormalities in heart, brain and skull

After demonstrating that TcMAC21 mice do not possess obvious mosaicism and that HSA21q genes show the expected expression patterns, we assessed phenotypes that occur frequently in DS and other DS mouse models.

#### Congenital heart defect (CHD)

CHDs are present in nearly 50% of people with DS and include septal defects, such as atrioventricular septal defects (AVSD), ventricular septal defects (VSD), and atrial septal defects (ASD), and outflow tract abnormalities, such as double outlet right ventricle (DORV). Several DS mouse models possess high rates of CHD (29–32). We examined hearts of TcMAC21 mice in a mixed, outbred background by both wet dissection at E18.5 and histology at E14.5. One TcMAC21 heart had both VSD and DORV in wet dissection (Fig. 4A), while another mouse had AVSD with a small dorsal mesenchymal protrusion (DMP) (Fig. 4B). We found that 28.6% of TcMAC21 had a structural defect of the heart by consolidating data of both assays. In TcMAC21, VSD was the predominant malformation, accounting for 21.4%, and AVSD accounting for 2.4%. The 24% frequency of septal defects in TcMAC21 is substantially greater than the 4% observed in Ts65Dn mice but smaller than the 38-55% reported in Tc1 (17, 29, 32, 33). AVSD occurs in 20% of babies with DS but is rare or absent from DS mouse models except in the presence of additional genetic modifiers (32, 34).

**Fig. 4.**
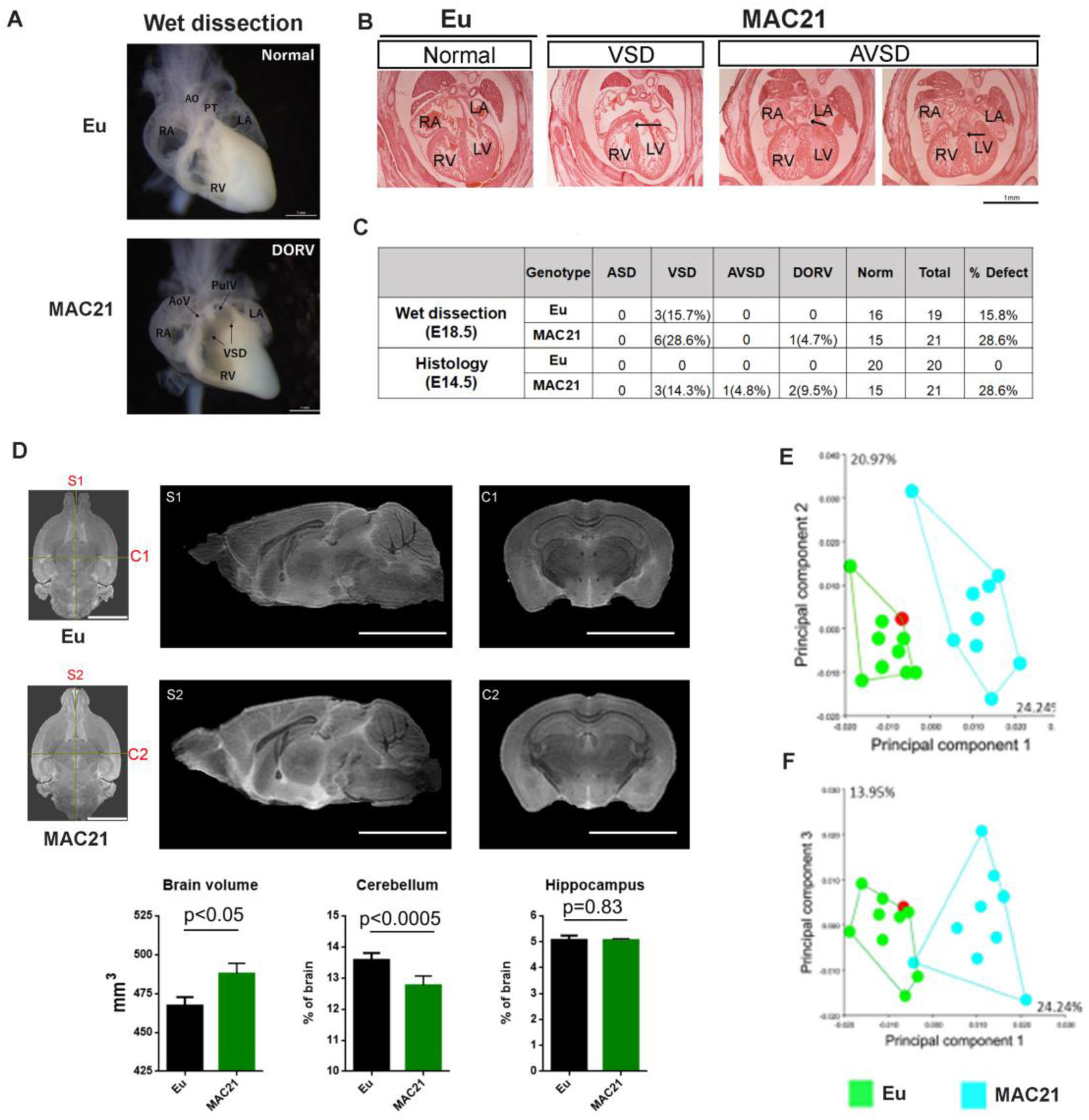
Morphological analysis of the heart, brain and skull in TcMAC21. (**A-C**) CHD analysis: (A) wet dissection of hearts from E18.5 Eu (top, normal) and TcMAC21(bottom, DORV and VSD (black arrow)), and a small slit-like conus VSD was also seen in TcMAC21; (B) histology of E14.5 Eu (normal) and two TcMAC21 mice with different heart defects, VSD and AVSD (black arrow indicating locations of defects); AO, aorta; AoV, aortic valve; LA, left atrium; LV, left ventricle; PT, pulmonary trunk; PulV, pulmonary valve; RA, right atrium; RV, right ventricle. (C) summary data of CHD analysis. (D) Representative T2-weighted MR images (top, *ex vivo*) and statistical analysis of whole brain volume, percentage of cerebellum and hippocampus relative to brain; mid-sagittal slices from Eu (S1) and TcMAC21 (S2) and coronal slices from Eu (C1) and TcMAC21 (C2), scale bar (5 mm); n=7 per group and data are analyzed by two-way ANOVA and expressed as mean ± SEM. (**E and F**) Results of a principal components analysis (PCA) of 3D geometric morphometric analysis of Eu and TcMAC21 cranial shape. PC1 shows a 24.2% separation between Eu and TcMAC21 mice while PC2 captures intraspecific variation (Eu (n=10) and TcMAC21 (n=9)).

#### Brain morphometric phenotypes

Cerebellar hypoplasia is among the few phenotypes present in every person with DS (35). Analysis of Ts65Dn identified a deficit in cerebellar granule cells (GC) and correctly predicted its occurrence in people with DS (36). This deficit is present in several DS mouse models as well (37, 38). Ventricle enlargement, which reportedly is associated with neurodegenerative diseases and aneuploidies including DS, was found in previous DS mouse models including Dp(16)1Yey, Ts1Cje and Ts2Cje (39, 40). Because ventricle collapse and cerebrospinal fluid (CSF) loss during brain fixation would cause false measurements in MRI (**Fig. S3A**), 3 out of 7 pairs were imaged live. TcMAC21 had larger total brain volume than Eu mice (487.9 ± 17.6 mm^3^ vs. 467.3 ± 14.6 mm^3^, p<0.05) (Fig. 4D). Nineteen brain subregions were further analyzed with absolute and percentage volume changes as outcome variables. Absolute volumes of some structures were significantly increased in TcMAC21, including lateral ventricle (p<0.003), thalamus (p<0.04), hypothalamus (p<0.02), superior colliculus (p<0.0001), amygdala (p<0.03) and nucleus accumbens (p=0.01) (**Table S2 and Fig. S3B**). Percentage volume of lateral ventricles (p=0.005) and superior colliculus (p<0.0001) were still significantly enlarged in TcMAC21 when normalized to total brain size (**Table S3**). Cerebellum was one of the few structures that had smaller absolute volume and the only structure whose percentage volume was significantly decreased in TcMAC21 (p<0.0005).

#### Craniofacial skeleton

The remarkable parallels of the impact of trisomy on skull development in individuals with DS and a number of mouse models is well-documented (41, 42). We used 3D geometric morphometric analysis to compare TcMAC21 and Eu cranial shape based on micro-CT images. Principal component analysis (PCA) showed separation between the two groups (Fig. 4E, F). In aspects of facial shape change, TcMAC21 mice have relatively more antero-posteriorly retracted and medio-laterally expanded (i.e., short and wide) snouts than do Eu mice. TcMAC21 neurocranium is slightly more “globular” than that of Eu. These differences are similar to those seen in other DS mouse models, such as Ts65Dn and Dp(16)1Yey (43, 44).

### TcMAC21 has various molecular and cellular pathological phenotypes

#### Higher APP protein levels without amyloid plaques

Individuals with trisomy 21 invariably display the plaques and tangles that represent two major pathologic characteristics of Alzheimer’s disease at an early age. These changes are linked to an extra copy of the amyloid precursor protein gene (*APP*) on HSA21 (45). *APP* was among the most highly expressed HSA21q genes in TcMAC21, and total APP mRNA (HSA21q + its mouse ortholog) in P1 forebrain of TcMAC21 was about twice as high as that of Eu **(Data file S2)**. In 15-24-month-old mice, total APP protein levels (HSA21q + its mouse ortholog) in hippocampus and cortex of TcMAC21 were significantly higher than Eu (p< 0.001, Fig. 5A). Both total Aβ40 and Aβ42 levels in brain were significantly increased in TcMAC21, but the Aβ40/Aβ42 ratio was not significantly different from Eu (Fig. 5B). We also found that total plasma Aβ40 levels in TcMAC21 at 7.5, 15 and 18 weeks of age were 2, 4 and 12 times as high as Eu, respectively **(Fig. S3C).** Despite elevated APP levels, TcMAC21, like other mouse models of DS, did not show spontaneous amyloid plaque formation at 15-24-month-old (Fig. 5C).

**Fig. 5.**
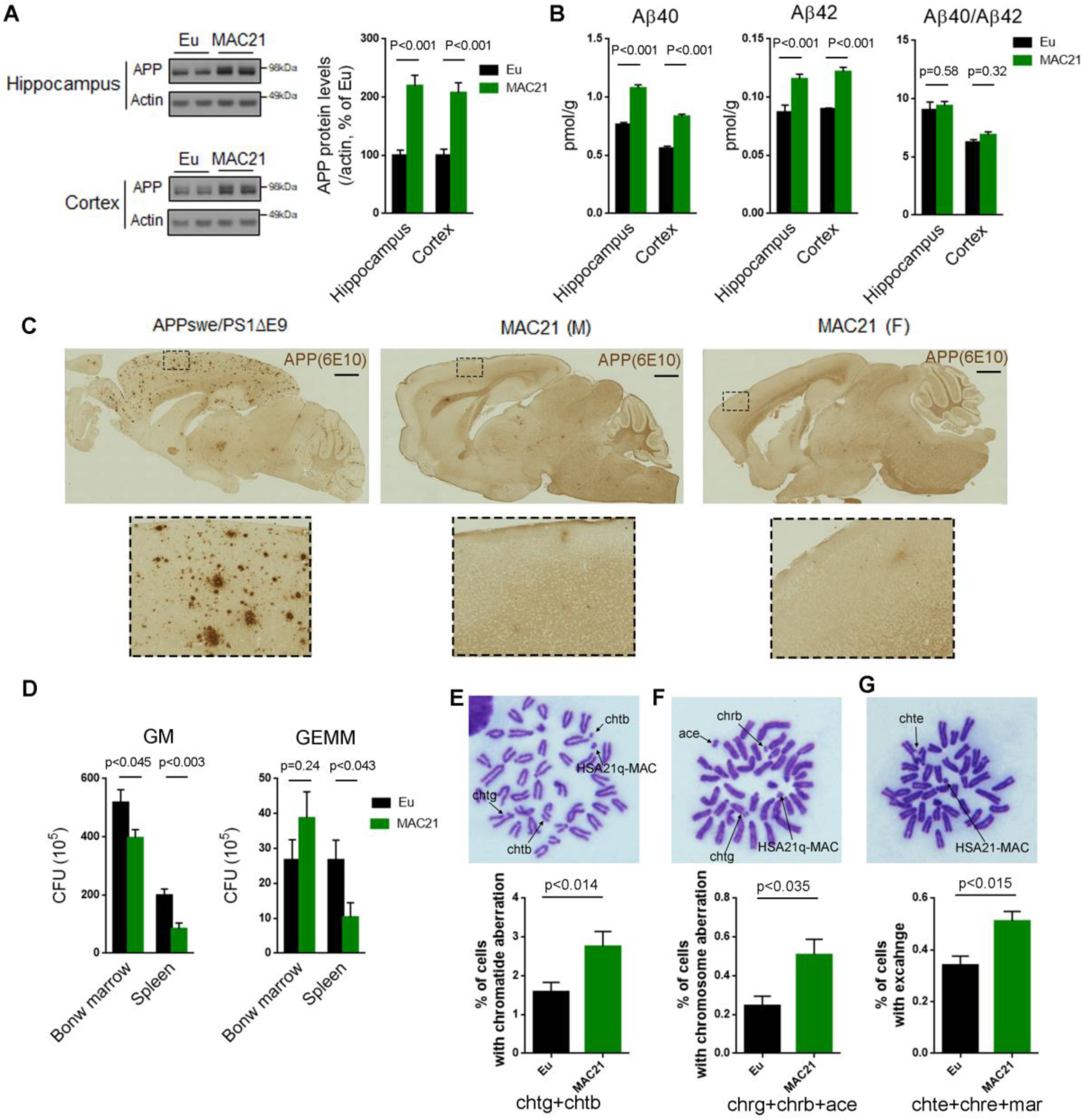
Evaluation of DS-like pathology in TcMAC21. (**A-C**) APP protein levels and amyloid plaques in brains of 15-24-month-old TcMAC21 and Eu (n=6 per group): (A) western blot of total APP in hippocampus and cortex; (B) ELISA of total Aβ40 and Aβ42 levels in hippocampus and cortex; (C) amyloid plaques visualized by immunostaining with APP antibody 6E10; *APPswe/PS1ΔE9* served as a positive control for plaque formation, scale bar (1 mm). (**D**) CFU level of GM and GEMM in TcMAC21 spleen and bone marrow (n=6 per group). (**E-G**) Chromatid and chromosome aberrations in bone marrow cells after X-ray irradiation (n=3 per group), (E) chromatid aberration, (F) chromosome aberration and (G) chromatid and/or chromosome exchange. Data are analyzed by two-tailed t-test and expressed as mean ± SEM.

#### Hematological abnormalities

People with DS often develop various hematopoietic disorders at different ages (46). In peripheral blood analyses, platelet counts were significantly increased in TcMAC21 compared to Eu (112.3 ± 7.8 versus 101.2 ± 6.8; p < 0.033) **(Table S4)**. TcMAC21 exhibited splenomegaly and white pulp hypertrophy by histology **(Fig. S4A)**. TcMAC21 showed a significantly reduced frequency of granulocyte/macrophage (GM) in bone marrow and reduced colony formation of GM and granulocyte/erythroid/monocyte/megakaryocyte (GEMM) in spleens using *in vitro* colony-forming assays (Fig. 5D).

#### Higher chromosomal radiosensitivity of bone marrow cells

Individuals with DS are reported to have increased sensitivity to X-rays and various chemical compounds that cause DNA damage in cultured lymphocytes (47). Radiation-induced suppression of the clonogenic activity of hematopoietic stem cells is associated with increased DNA double strand breaks (48). We observed that following X-ray irradiation, bone marrow cells from TcMAC21 had a significantly higher frequency of aberrations and exchanges in chromatids and chromosomes than Eu (Fig. 5E-G).

#### TcMAC21 has fine breeding but significant growth defect

Four BDF1.TcMAC21 females crossed to B6 males produced 15 litters comprising 90 pups on F1 background. Litter size ranged from 2 to 9 with an average of 6 pups; 48% of offspring were TcMAC21 of which 54% were male (**Table S5**). Thus, TcMAC21 husbandry appears to be the same or somewhat better than that of Ts65Dn, which can produce ample numbers of offspring for most experiments with a reasonably sized colony.

Individuals with DS show a slower growth rate with delayed developmental milestones, which is reflected in several mouse models of DS (49). We measured the mass of TcMAC21 and Eu mice from P1 through P90. The average mass for the TcMAC21 cohort was consistently smaller than Eu (Fig. 6A **and Data file 3)**. On average, adult TcMAC21 mice were about 25% smaller than Eu at P90. Both males and females weighed less than their Eu counterparts (Fig. 6B).

**Fig. 6.**
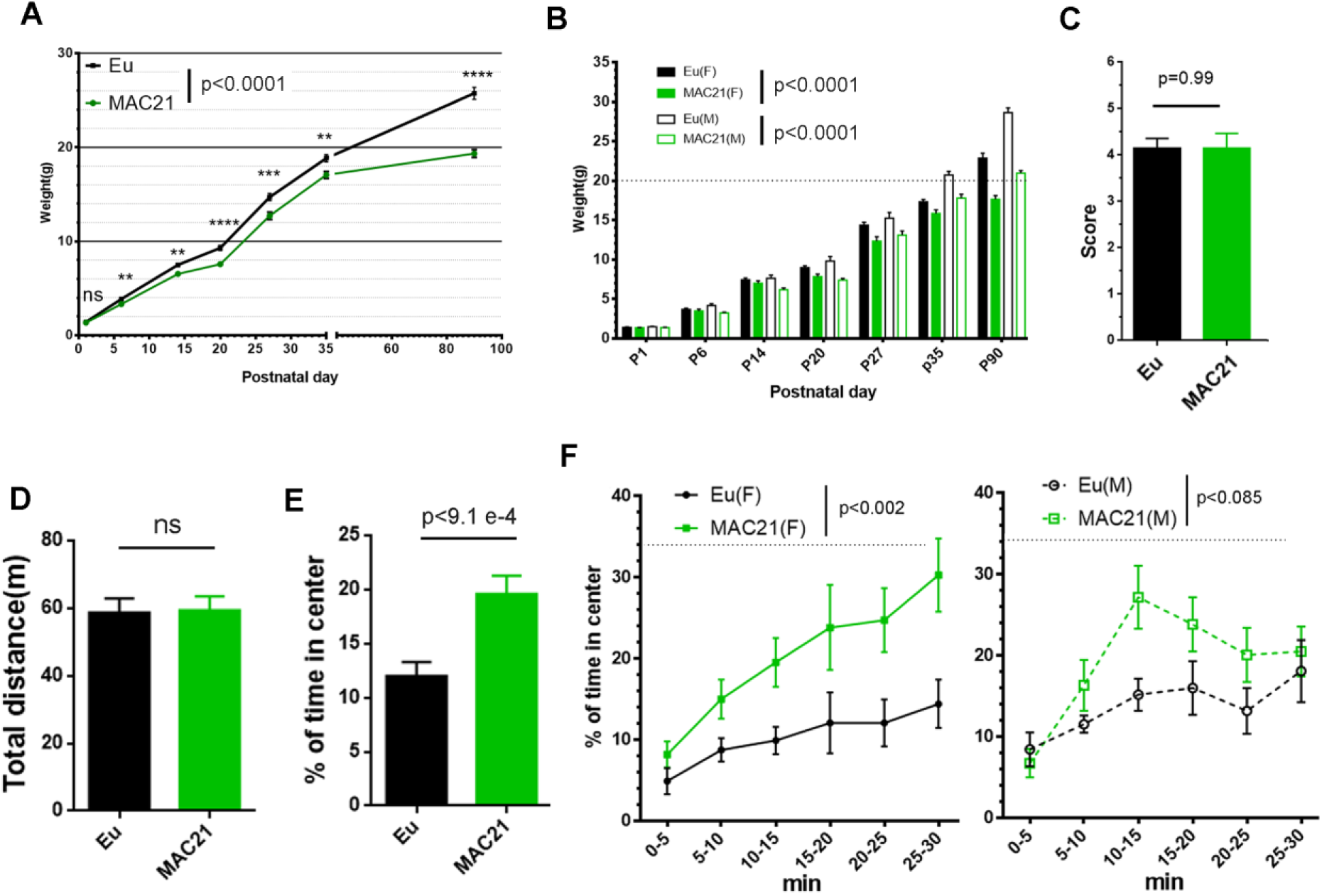
Growth profile, nesting and open field in TcMAC21. (**A-B**) The mass of TcMAC21 and Eu were measured during postnatal development at P1, P6, P14, P20, P27, P35 and P90: (A) female and male were consolidated (n≥23 per group; the overall genetic effect is analyzed using two-way ANOVA; genetic effect at each timepoint is analyzed using two-tailed t-test, *P < 0.05, **P < 0.01, ***P < 0.001, ****P < 0.0001); (B) male TcMAC21 are larger than females (two-way ANOVA and Tukey’s post-hoc). (**C**) Nesting (female: Eu (n=19), TcMAC21 (n=13); male: Eu (n=7), TcMAC21 (n=7); two-tailed t-test). (**D-F**) 30 min open field: (D) total travel distance, (E) percent of the time spent in central zone and (F) dynamics of time spent in the central zone (female: Eu (n=12), TcMAC21 (n=11); male: Eu (n=11), TcMAC21 (n=12); female and male were consolidated and two-tailed t-test used in (D and E); female and male were significantly different (repeated measures ANOVA with LSD-test). Data are expressed as mean ± SEM and see also Data file S3 and Data file S4.

### TcMAC21 has significant learning and memory defects

#### TcMAC21 displays normal nest-building activity

Nest construction is a complex task that may require the coordination of various parts of brain including hippocampus, prefrontal cortex, and cerebellum (50–52). Ts65Dn mice show defects in nesting (38, 53). However, adult TcMAC21 nesting behavior was equivalent to Eu (Fig. 6C).

#### TcMAC21 shows reduced anxiety in open field

We used a 30 min novel open field paradigm to assess novelty-induced exploratory activity and anxiety. TcMAC21 exploratory activity was not different from Eu based on total distance traveled (Fig. 6D), in contrast to hyperactivity in Ts65Dn (54). Both male and female TcMAC21 spent significantly more time exploring the center area compared to Eu, indicating less anxiety in TcMAC21 (Fig. 6, E and F).

#### TcMAC21 shows deficits in Morris water maze (MWM)

The visual discrimination task showed no significant difference between TcMAC21 and Eu, indicating that TcMAC21 has normal visual ability and goal-directed behavior **(Fig. S5, C and D)**. There was no difference in swimming speed between TcMAC21 and Eu, but males of both genotypes swam faster than the females of corresponding genotype **(Fig. S6D-I)**. To avoid possible group-specific differences in swimming speed, acquisition trials are reported as escape distance rather than latency. Throughout four days of MWM training, both TcMAC21 and Eu reduced escape distance to find the hidden target platform in acquisition trials, but the improvement was significantly slower in TcMAC21 than Eu (p<0.015, Fig. 7A). To test spatial learning and memory, both short delay (30 min) and long delay (24h and 72h) probe trials were performed (Fig. 7C-F). In the short delay probe trials, both TcMAC21 and Eu increased time spent in the target area from day 1 to day 4, but TcMAC21 performance lagged significantly behind that of Eu throughout the training days (p< 0.002, Fig. 7C). Spatial memory disruption in TcMAC21 was more significant in long delay probe trials **(p<9.1e-5**, Fig. 7E). Poor MWM performance occurred in both female and male TcMAC21.

**Fig. 7.**
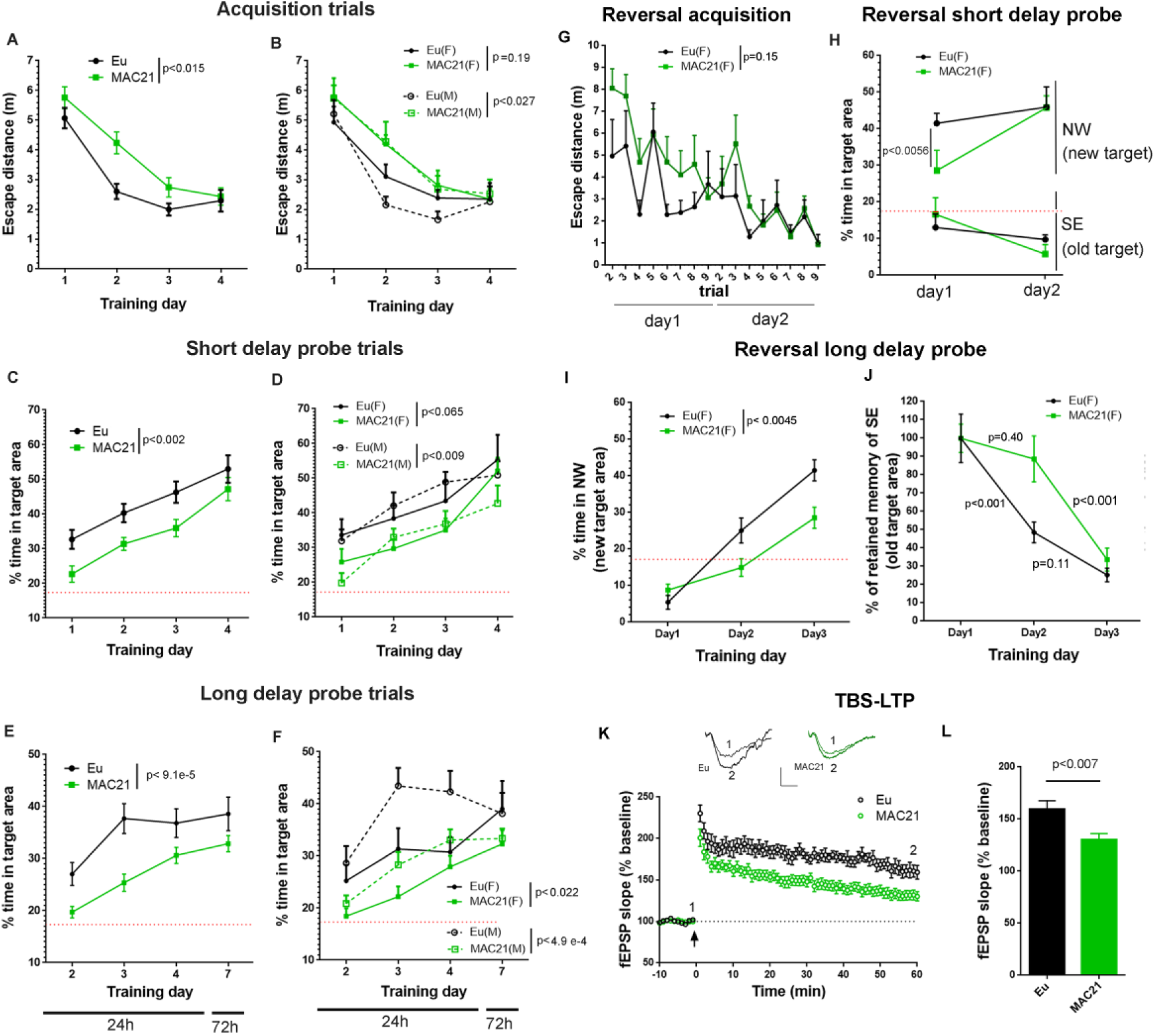
Learning and memory deficits in TcMAC21. **(A-F)** Classic MWM (female: Eu (n=10), TcMAC21 (n=11); male: Eu (n=11), TcMAC21 (n=12); female and male data are consolidated in (A, C and E) and separated in (B, D and F)): (A and B) acquisition trials (average distance of 8 trials per day); (C and D) short delay probe trials; (E and F) long delay probe trials. **(G-J)** RRWM (female: Eu (n=8), TcMAC21 (n=9)): (G) acquisition trials in RRWM1; (H) short delay probe trials in RRWM1; long delay probe trials from day 1 to day 3 for comparing abilities of learning and memorizing a new target (I) and inhibiting memory of previous target (J). **(K)** TBS-induced LTP at SC-CA1 synapses in Eu and TcMAC21. Normalized fEPSP slopes are plotted every 1 min. Sample traces represent fEPSPs taken before “1” and 60 min after TBS stimulation “2”. Arrow indicates LTP induction, and scale bars represent 0.5 mV (vertical), 5 ms (horizontal). **(L)** The amplitude of fEPSP slopes are averaged at 60 min after the stimulation (Eu (n=5, 13 slices), MAC21 (n=5, 14 slices)). Repeated measures ANOVA with LSD-post hoc test was used in (A-J) and two-tailed t-test was used in (K and L); data are expressed as mean ± SEM and see also Data file S4.

#### TcMAC21 has impairment in repetitive reversal water maze (RRWM)

To study whether the abilities to learn a new platform position and inhibit old memory of previous platform position were intact in TcMAC21, we tested female mice in RRWM immediately after MWM (**Fig. S7A**). In reversal 1, TcMAC21 performed relatively worse than Eu in acquisition trials (p=0.15, Fig. 7G). In short delay probe trials, TcMAC21 was significantly impaired in remembering the new target platform on day 1 (p<0.006, Fig. 7H) but caught up to Eu on day 2. The deficit in remembering the new platform was more obvious in long delay (24h) probe trials (p<0.005, Fig. 7I). Twenty-four hours after reversal day 1 training of the new platform, average memory retention of the old platform was >85% in TcMAC21 and >50% in Eu (Fig. 7J), indicating that TcMAC21 had difficulty in suppressing irrelevant information.

#### TcMAC21 has reduced hippocampal long-term potentiation (LTP)

In numerous models of human disease, decreased LTP is frequently correlated with diminished learning and memory. We assessed synaptic plasticity in TcMAC21 by theta burst stimulation (TBS)-induced LTP at SC-CA1 synapses on hippocampal slices. TcMAC21 had a significant reduction in LTP compared to Eu littermates (Fig. 7K-L). Together, these results showed that TcMAC21 had significant deficits in learning, memory, and synaptic plasticity.

## DISCUSSION

DS research has made impressive gains in the last 25 years, substantially fueled by the development of trisomic mouse models. Before the advent of segmental trisomy models, assessments of gene dosage effects in development and function were often based on single gene transgenic mice and necessarily proceeded on a gene by gene basis. Results were interpreted in the context of the reductionist assumption that the clinical presentation is nothing more than the sum of additive, independent effects of (a subset of) HSA21 genes (55). A considerably more nuanced view has emerged with the use of trisomic models, showing interactions of HSA21 genes, a transcriptome in which disomic as well as trisomic gene expression is perturbed throughout the genome, and genetic interaction of trisomic genes whose elevated expression potentiates phenotypes that require disomic modifiers (56). Despite this progress, fundamental questions remain about the most complex genetic insult compatible with survival beyond term.

In this study, we applied MMGT system and MMCT to establish a new DS mouse model, TcMAC21, which contains a nearly complete, freely segregating HSA21q as a mouse artificial chromosome. These mice have no detectable mosaicism in a broad spectrum of tissues and cell types. TcMAC21 is the most complete genetic model of DS created to date with 93% of HSA21q PCGs and about 79% of non-PCGs present. Representative human genes are transcribed in a tissue appropriate manner and the total transcript increase (human plus mouse ortholog) is broadly at the level that is expected for an additional copy of the affected genes. These changes in gene expression levels are correlated with several phenotypes that occur in people with trisomy 21 and in other mouse models of DS, including small size (short stature); dysmorphology of the brain including cerebellar hypoplasia; anomalies of the craniofacial skeleton; congenital heart defects; elevated expression of the Alzheimer-related *APP* gene and its cleavage products; sensitivity of chromosomes to radiation-induced aberrations; and deficits in learning and memory and LTP, their physiological correlate.

The TcMAC21 model provides the best current representation of gene dosage comparable to that of DS, and thus should prove valuable as a new model for preclinical studies aimed at ameliorating effects of trisomy 21 and maximizing opportunities for individuals with trisomy 21 to develop their full potentials. Moreover, the successful experience in generating and characterizing the first rodent model with a hybrid whole human chromosome paves the path for generating other hybrid chromosome animal models for other human aneuploidies.

## Materials and Methods

### Animals

All procedures related to animal care and treatment were approved by each local University/Institutional Animal Care and Use Committee and met the guidelines of the National Institute of Health and RIKEN Guide for the Care and Use of Laboratory Animals. TcMAC21 mice were initially maintained on an outbred background (ICR strain mice). After crossing for 8 generations onto BDF1 (C57BL/6J (B6) × DBA/2J (D2)), TcMAC21 were transferred to the Riken Animal Resource (BRC No. RBRC05796 and STOCK Tc (HSA21q-MAC1)). Except as noted otherwise, phenotypic characterizations of TcMAC21, including histology of CHD, MRI, craniofacial morphology, husbandry, growth profile and behavioral tests, were performed using mice with 75% B6/25% DBA on average.

### Chromosome engineering to produce HSA21q-MAC

The HSA21q-MAC was constructed using a previously described Mb-sized gene cloning system with the MAC1 vector (57). A loxP site was inserted at position 13021348-13028858, NC_000021.9 in a Chr.21 in DT40 cells as described previously (27). The modified hChr.21 (hChr.21-loxP) was transferred to CHO (MAC1) cells via MMCT as described previously (21). MAC1 contains a centromere from mouse chromosome 11, EGFP flanked by HS4 insulators, PGK-neo, loxP-3′HPRT site, PGK-puro, and telomeres (22). Hprt-deficient Chinese hamster ovary (CHO hprt−/−) cells containing MAC1 were maintained in Ham’s F-12 nutrient mixture. Cre-recombinase expression vectors were transfected into CHO cells containing the MAC1 and the HSA21-loxP using Lipofectamine 2000. The cell culture and colony expansion were performed as described previously (58). The site-specific reciprocal translocation between the MAC1 and the HSA21-loxP was confirmed by PCR (**Table S6**) and FISH analyses. Mouse ES cells were fused with microcells prepared from the donor hybrid cells and selected with G418. Parental mouse ES cell line, TT2F, and derivatives were maintained on the mitomycin-treated mouse embryonic fibroblasts.

### FISH and G-banding

FISH was performed using fixed metaphase or interphase spreads. Slides were hybridized with digoxigenin-labeled (Roche, Basel, Switzerland) human COT-1 DNA (Invitrogen) and biotin-labeled mouse COT-1 or minor satellite DNA (a gift from Dr. Vladimir from NIH), essentially as described previously (21, 59). For Giemsa banding, chromosome spreads were prepared and stained as described (60).

### Whole Genome and RNA Sequencing

#### WGS

DNA from a TcMAC21 mouse was purified and sequenced for four different runs using the Illumina NextSeq 500 DNA sequencer (**Data file S1**). After cleaning, the short reads were mapped to whole genome sequences of mouse (GRCm38) and human chromosome 21 (NCBI NC_000021.9) using CLC Genomics Workbench ver. 9.5. A total of 974.89 million reads were mapped to the reference sequences. Among these reads, 4.72 million reads were mapped to HSA21, GRCh38.p13 Primary Assembly. Most of the mapped reads were located between positions 13.0M and 46.7M. The effective depth of coverage was 25.5×. All raw read data of WGS were deposited to DDBJ Sequence Read Archive (DRA) under accession number DRA008337 and DRA008342. ***RNA-Seq:*** RNA was extracted from forebrains of Eu and TcMAC21 at P1. Standard mRNA purification and library preparation were conducted using NEBNext® Poly (A) mRNA Magnetic Isolation (E7490) and NEBNext® Ultra™ II RNA Library Prep Kit (E7770). Library quality was assessed via Agilent 2100 Bioanalyzer for DNA High sensitivity DNA chip. The prepared library was sequenced using HiSeq2500 Flowcell with 100 bp paired-end reads, with each sample containing approximately 50-60 million reads. Sequence was assessed with fastqc, and 30bp were trimmed from each sequence. HSA21 reference was extracted and appended onto the whole mouse genome reference sequence to create the modified reference. Reads were then aligned with TopHat2. Sim4 and Leaff were used for cross-species analysis. Standard DEseq methodology was used for differential gene expression analysis.

### CHD analysis by wet dissection and histology

#### Wet dissection

E18.5 mouse fetuses were removed and sacrificed, and hearts were flushed with PBS via the umbilical vein and then fixed in 4% PFA. The hearts were examined for cardiovascular anomalies under a dissecting microscope. ***Histology:*** E14.5 embryos were collected and fixed in 10% formalin for 48h. Tissues were embedded in paraffin, sectioned at 10 uM, and stained with hematoxylin/eosin. The heart was analyzed via dissection stereomicroscope (Nikon SMZ1500, Japan).

### Brain morphometry by MRI

4.5-month-old TcMAC21 and Eu mice were used for either *ex vivo* or *in vivo* MRI (61). For the *ex vivo* scan, 4 pairs of mice were perfused with 4% PFA after PBS and heads were post-fixed for 1 week, then kept in PBS for 3 days. Heads were stored in Fomblin to prevent dehydration during imaging in an 11.7 Tesla scanner (vertical bore, Bruker Biospin, Billerica, MA). 3D T2-weighted images were acquired on an 11.7 Tesla Bruker scanner (Bruker Biospin, Billerica, MA, USA) with the resolution = 0.08 mm × 0.08 mm × 0.08 mm. For the *in vivo* scan, 3 pairs of mice were anesthetized with isoflurane and monitored with a physiological monitoring system during imaging in 9.4 Tesla scanner (Bruker Biospin, Billerica, MA, USA). 3D T2-weighted images were acquired at resolution = 0.1 mm × 0.1 mm × 0.25 mm. For analysis, both *ex vivo* and *in vivo* images were first aligned to the template image using automated image registration software (Diffeomap, www.mristudio.org) and adjusted to an isotropic resolution of 0.0625 mm × 0.0625 mm × 0.0625mm.

### Craniofacial morphology by micro-CT

Nine 3D anatomical facial landmarks were collected on micro-CT of 9 TcMAC21 and 10 Eu at age of 4.5 month old. Each specimen’s landmark configuration was superimposed using the generalized Procrustes analysis (GPA). This method extracts shape coordinates from the original specimen landmark configurations by translating, scaling, and rotating the data and subsequently yields a measure for size called centroid size (Dryden Mardia, 1992). To examine cranial shape variation in the sample we used principal component analysis (PCA). All analyses were conducted in MorphoJ and software R.

### Western blot, ELISA, and amyloid plaque staining

15-24-month-old Eu and TcMAC21 mice (n = 6 each) were perfused with PBS, then a half brain was used to make lysate for Western blot and ELISA, and the other half was fixed in 4% PFA. ***Western Blot and ELISA:*** cortex and hippocampus were removed and lysed with RIPA buffer. Protein extracts were separated by 4-12% SDS-PAGE, transferred to PVDF membranes and then probed with APP antibody (Millipore, MAB348) and beta actin. For ELISA, the above lysate was spun at 16,000g for 30 min at 4°C. Supernatant was analyzed to determine Aβ levels by human/rat β amyloid ELISA kit from Wako (Aβ40: Cat# 294-64701; Aβ42: Cat# 290-62601). ***Amyloid plaque staining:*** mouse brains were fixed by immersion in 10% PFA, embedded in paraffin, and sectioned at 5 µm. Sections were deparaffinized and protein antigenicity was unmasked, and then the endogenous peroxidase activity was inhibited with 1.5% hydrogen peroxide. Sections were incubated with mouse anti-β-amyloid (Covance, Cat# SIG-39320), biotinylated goat anti-mouse IgG, and Avidin/Biotin mixture and then developed in 3,3′-Diaminobenzidine (DAB).

### Peripheral blood analyses

10-12-month-old mice were bled from the retro-orbital plexus under general anesthesia and euthanized for further analyses. Hematopoietic indices were measured with a hemocytometer (Nihon Koden).

### Colony-forming cell assays

Bone marrow and splenic mononucleated cells were incubated in duplicate at cell concentrations of 2×10^4^ and 2×10^5^/mL, respectively, in MethoCult-M3434 semisolid culture medium (STEMCELL TECHNOLOGIES). Colonies were scored on day 3 for erythroid colony-forming units (CFU-Es) and on day 8 for granulocyte/erythroid/monocyte/megakaryocyte colony-forming units (CFU-GEMMs), granulocyte/monocyte colony-forming units (CFU-GMs), and monocyte colony-forming units (CFU-Ms). For the megakaryocyte colony-forming units (CFU-Megs) assays, 5×10^4^/mL bone marrow and 5×10^5^/mL spleen cells were cultured with MegaCult-C Kit (STEMCELL TECHNOLOGIES) for 9 days, and colonies were stained for acetylcholine esterase (AChE) in accordance with manufacturer’s recommendation.

### Radiosensitivity test

TcMAC21 and Eu littermates at 16-18 weeks old received whole body X-ray irradiation at the rate of 0.25 Gy/min for 12 min using MX-160Lab (mediXtec Japan Corporation). After 8h, irradiated mice were administrated colcemid (1.5mg/kg) (Demecolcine, Sigma-Aldrich, USA) by intraperitoneal injection. After 3h, bone marrow cells were harvested from tibia and femurs of both legs and then fixed in Carnoy’s solution. Chromosome spreads were prepared by dropping the cell suspension on a grass slide and stained with 5% Giemsa solution (Merck Millipore, Germany) for 15 min. The analyses for chromosome aberrations were performed using the stained metaphase spreads and classification of chromosome aberrations according to (62). For each mouse, >30 mitotic cells were analyzed, and aberration rate was expressed as % of total cell analyzed.

### Behavioral tests

Both TcMac21 and Eu were used for behavioral tests in the sequence, open field, visual discrimination water maze test, MWM, and RRWM. The ANY-maze tracking system (Stoelting Co.) was used to collect data. ***Open field:*** was performed after three days of handling in the same room consisting of indirect diffusing light (∼150 lux) for all animals. The whole arena size was 37 cm X 37 cm, and the center area (21.6 cm X 21.6 cm) was 34% of the whole arena. Distance traveled and percentage of time spent in center were analyzed in 30 min and 5-min-bins. One female mouse was excluded because of bad tracking: female: Eu (n=12), TcMAC21 (n=11); male: Eu (n=11), TcMAC21 (n=12). ***Visual discrimination (VD):*** the day before VD, all mice were pre-trained to climb and stay on a submerged platform (10 cm X 10 cm) in a small clear water pool (45 cm diameter) for five trials (63). Non-toxic white tempera paint was used to make the platform invisible in a water tank 126 cm in diameter **(Fig. S5A)**. There was no spatial cues, but the location of the platform was made visible by attaching a black extension, 4 cm above water surface. The platform position was changed every two trials from W to E to S for a total six trials **(Fig. S5B)**. ***Classic MWM:*** for all animals, the same spatial cues were used and MWM was performed for four days (each day had 10 trials, which included 8 acquisition trials plus 2 probe trials of short-(30 min) and long-delay (24h)). A longest delay probe trial was conducted 72h after the fourth training day (**Fig. S6A)**. The platform remained in the same position in “SE quadrant” during the MWM test, with the water temperature at 22 ± 2°C. The platform was hidden ∼1.8 cm below the water surface and 60 seconds was the maximum time allowed in acquisition trials, and if a mouse did not find the platform, the tester would visually or manually guide it to the platform. For the probe trials, the platform was lowered to a position that mice were not able to climb onto for 30-40s. At the end of probe trials, the collapsed platform was raised to the same position used in the acquisition trial and the tester guided the mouse to the platform, which helped maintain the same response-reinforcement contingency of the acquisition. If a mouse continually failed to follow the tester’s guidance to reach the platform, it was excluded from analysis. Three female mice were excluded by this standard, which left these mice (female: Eu (n=10), TcMAC21 (n=11); male: Eu (n=11), TcMAC21 (n=12)). ***RRWM:*** following the classic MWM, 9 TcMAC21 and 10 Eu female mice were tested in RRWM without changing any spatial cues. RRWM consisted of two reversal learning tests in which the platform was first relocated to NW from SE for two days and then relocated to SW for another two days **(Fig. S7A)**. Trial 1 of reversal day 1 was the same as the 72h delay probe trial in the classic MWM. Each day had 10 trials including 8 acquisition trials and 2 probe trials for short delay(30 min) and long delay (24h). Excluding 2 Eu female mice that failed to follow tester’s guidance, 9 TcMAC21 and 8 Eu female mice were analyzed for RRWM. ***Nesting:*** 3-4-month-old TcMAC21 mice and their littermates were singly housed with an intact compressed cotton nesting pad in a new cage for 24h. The mouse was removed, and the cage was photographed. An observer blinded to the genotypes of mice scored “nesting quality”: 1, <50% of nesting square shredded but not organized; 2, <50% of nesting square shredded and organized or 50-99% of nesting square shredded but not organized; 3, 50-99% of nesting square shredded and organized or 100% of nesting square shredded but not organized; 4, 100% of nesting square shredded and organized into a large nest that covers less than half of the area of the cage; 5, 100% of nesting square shredded and organized into a compact nest that covers less than a quarter of the area of the cage; or 6, 100% of nesting square shredded and organized into a small nest with rounded edges and a “donut hole” center. Both female (Eu (n=19), TcMAC21 (n=13)) and male (Eu (n=7), TcMAC21 (n=7)) mice were tested.

### Electrophysiology

Following behavioral tests, 5 pairs of Eu:TcMAC21 mice (3-4 months old) were deeply anesthetized with inhaled isoflurane and then perfused with ice-cold oxygenated cutting solution containing (in mM): 110 choline chloride, 7 MgCl2, 2.5 KCl, 0.5 CaCl2, 1.3 NaH2PO4, 25 NaHCO3, 20 glucose, saturated with 95% O2 and 5% CO2. Transverse hippocampal slices (400 mm) were cut using a vibratome (VT-1200S, Leica) and transferred to artificial cerebrospinal fluid (aCSF) containing (in mM): 125 NaCl, 2.5 KCl, 2.5 CaCl2, 1.3 MgCl2, 1.3 NaH2PO4, 25 NaHCO3, 10 glucose, saturated with 95% O2 and 5% CO2. Slices were allowed to recover for 40 min at 32 °C and then at room temperature for at least 2 h before recording. Picrotoxin (100 µM) was added to block inhibitory transmission. Slices were transferred to the recording chamber and perfused continuously with aCSF (flow rate at 2-3 mL/min) at room temperature. A cut between CA3 and CA1 was made to minimize recurrent activity during recording. A concentric bipolar electrode (World Precision Instruments) was placed in the middle of CA1 stratum radiatum to stimulate Schaffer collateral. Field EPSPs (fEPSPs) from the CA1 neurons were recorded with a glass pipette (2-3 MΩ) filled with aCSF. Constant current pulses (70-100 µA, 100 µs) were delivered at 0.033 Hz by a STG 400 stimulator. The stimulus intensity was adjusted to evoke 40%–50% of the maximal response. LTP was induced by theta burst stimulation (TBS) consisting of a single train of 5 bursts at 5 Hz, and each burst contained 4 pulses at 100 Hz. The recording and data analysis were performed by investigators blinded to mouse genotype.

## Acknowledgments

We thank Toko Kurosaki, Yukako Sumida, Hiromichi Kohno, Masami Morimura, Kei Yoshida, Eri Kaneda, Akiko Ashiba, Dr. Kazuomi Nakamura, Rina Ohnishi, Yuwna Yakura, Etsuya Ueno, Dr. Yuichi Iida, Dr. Yoshiteru Kai, Motoshi Kimura, Chie Ishihara, Kiyoko Kawakami, and Chiga Igawa at Tottori University for their technical assistance; as well as Dr. Hiroyuki Kugoh, Dr. Masaharu Hiratsuka, Dr. Tetsuya Ohbayashi, Dr. Hiroyuki Satofuka, and Dr. Takahito Ohira at Tottori University, Dr. Oomiya Yoshihiro at Health Research Institute, National Institute of Advanced Industrial Science and Technology (AIST), and Dr. Shigeharu Wakana, Dr. Tamio Furuse, Dr. Ikuko Yamada at Technology and Development Team for Mouse Phenotype Analysis Japan Mouse Clinic, RIKEN BioResource Center (BRC) for critical discussions. This research was partly performed at the Tottori Bio Frontier managed by Tottori prefecture.

