## Supplementary Information for "A non-mosaic humanized mouse model of Down syndrome, trisomy of a nearly complete long arm of human chromosome 21 in mouse chromosome background"

**This PDF file includes:**

Figures S1 to S7
Tables S1 to S6

**Other supplementary materials for this manuscript include the following:**

Datasets S1 to S4

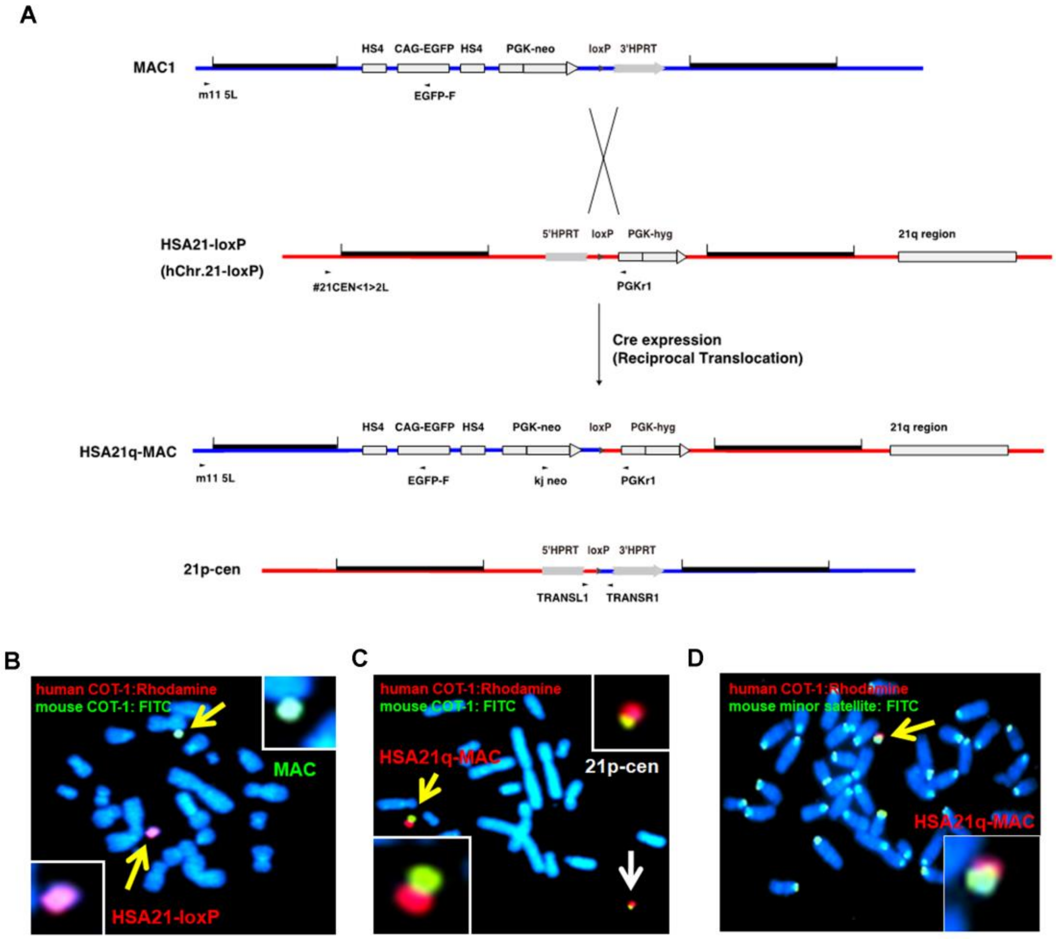


**Fig. S1.** (**A**) Information for vectors used to generate HSA21q-MAC. (**B-D**) FISH images to verify the critical steps: (B) before Cre-loxP mediated recombination, CHO cells contained a modified hChr.21 and a MAC; (C) after recombination, CHO cells contained the HSA21q-MAC (yellow arrow) and by-product (21p-cen) (white arrow); (D) mouse ES cell containing the HSA21q-MAC (yellow arrow).

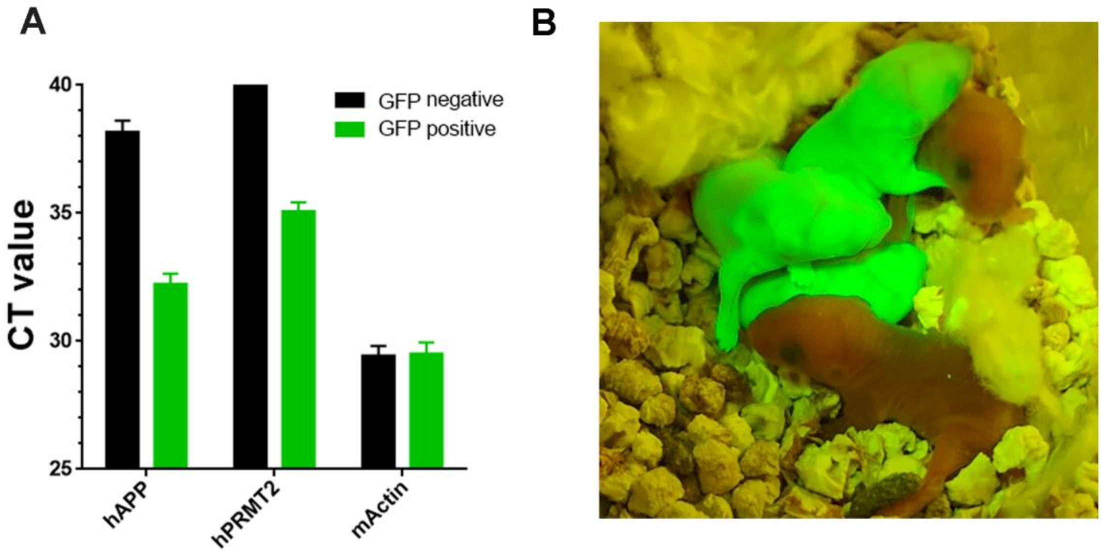


**Fig. S2. GFP is a reliable marker for genotyping MAC21.** (**A**) Tails from GFP negative and positive TcMAC21 littermates were analyzed using Taqman RT-PCR assay for testing expression of human-APP (hAPP, centromere-proximal), human-PRMT2 (hPRMT2, telomere-proximal), and mouse actin (mActin, endogenous control). N=12 per group. (**B**) TcMAC21 and Eu pups were visualized with GFP flashlights.

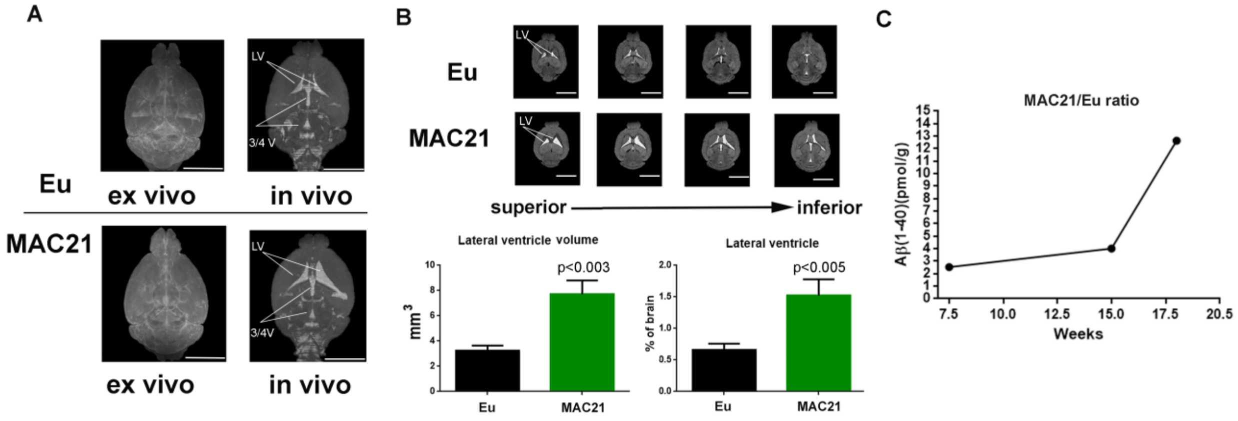


**Fig. S3.** (**A**) 3D rendering of Eu and TcMAC21 brains MRI images. No clear ventricle space was seen in fixed samples (*ex vivo* MRI) and clear ventricles and CSF were seen in live samples (*in vivo* MRI). LV: lateral ventricle, 3/4 V: the third and fourth ventricle; scale bar (5 mm) (**B**) LV volume of live Eu and TcMAC21 mice was analyzed. LV was visualized in 4 matching transverse levels of MRI images beginning at the top and working to the bottom of LV (adjacent levels spaced 312.5um apart, scale bar (5 mm)). Absolute and percentage of brain volumes of LV were compared (N=3 pairs and data are analyzed by two-tailed t-test and expressed as mean ± SEM). (**C**) ELISA assay of total Aβ(1-40) in plasma from Eu and TcMAC21 littermates at different ages. For each timepoint, 1 pair of littermates were analyzed.

A

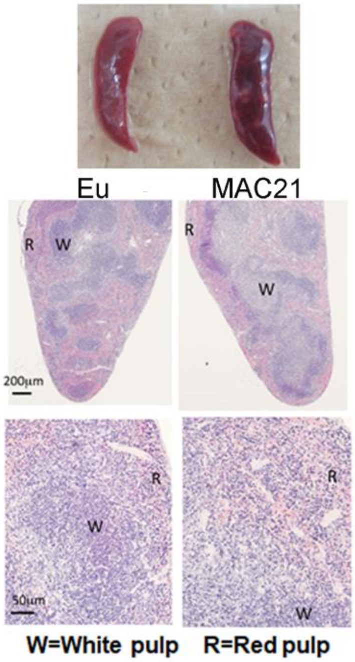


B

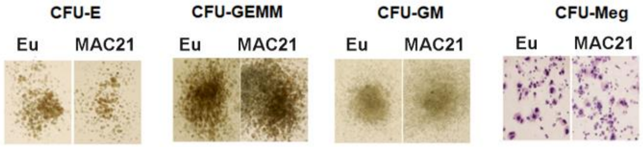


C

| Bone Marrow | (n) | CFU-E ($/10^5$) | CFU-GEMM ($/10^5$) | CFU-GM ($/10^5$) | CFU-M ($/10^5$) | CFU-Meg ($/10^6$) |
|---|---|---|---|---|---|---|
| Eu | 6 | 242 ± 52 | 26.7 ± 14.2 | 518 ± 109 | 29.3 ± 13.6 | 109 ± 62 |
| MAC21 | 6 | 209 ± 43 | 38.7 ± 18.5 | 396 ± 71 | 53.0 ± 60.7 | 140 ± 42 n=5 |
| p-value (T-test) | | $2.48 \times 10^{-1}$ | $2.36 \times 10^{-1}$ | **$4.47 \times 10^{-2}$** | $3.90 \times 10^{-1}$ | $3.72 \times 10^{-1}$ |

D

| Spleen | (n) | CFU-E ($/10^5$) | CFU-GEMM ($/10^5$) | CFU-GM ($/10^5$) | CFU-M ($/10^5$) | CFU-Meg ($/10^6$) |
|---|---|---|---|---|---|---|
| Eu | 6 | 144 ± 43 | 26.7 ± 13.9 | 199 ± 53 | 17.7 ± 8.8 | 167 ± 74 n=5 |
| MAC21 | 6 | 145 ± 87 | 10.3 ± 10.2 | 84 ± 50 | 24.7 ± 22.8 | 153 ± 67 |
| p-value (T-test) | | $9.94 \times 10^{-1}$ | **$4.24 \times 10^{-2}$** | **$3.42 \times 10^{-3}$** | $4.99 \times 10^{-1}$ | $7.46 \times 10^{-1}$ |

**Fig. S4.** (**A**) Histological analysis of spleen (R and W indicate red pulp and white pulp, respectively). (**B-D**) Colony assay of spleen and bone marrow: (B) representative colony images of bone marrow; (C and D) statistical analysis of colony-formation unites (CFU) of erythroid (E) granulocyte/erythroid/monocyte/megakaryocyte (GEMM), granulocyte/macrophage (GM), macrophage (M) and megakaryocyte (Meg) in bone marrow (C) and spleen (D), n=6 per group unless otherwise stated in figs, and data are analyzed by two-tailed t-test and expressed as mean ± SD.

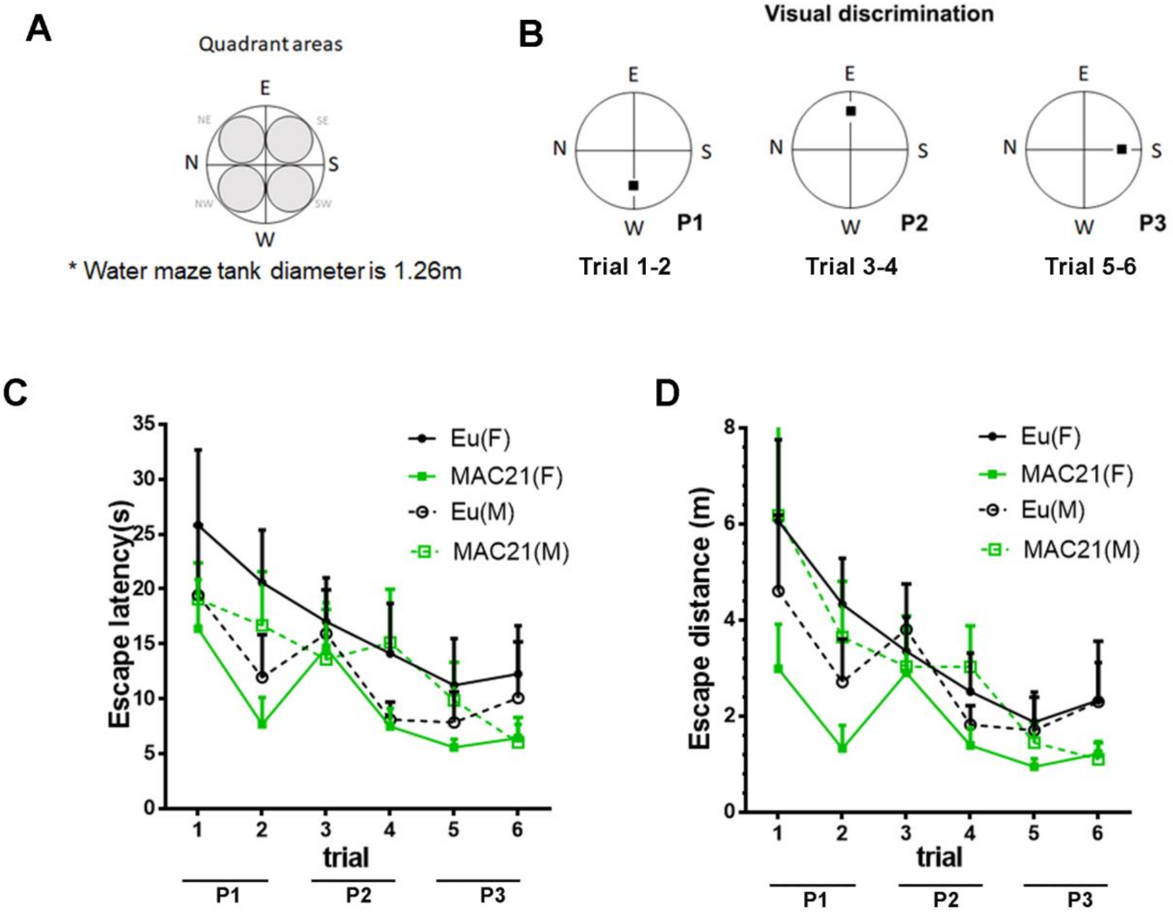


**Fig. S5. Visual discrimination in water maze.** (**A**) Water tank specifications. (**B**) Design of visual discrimination: platform changes every two trials from W, to E, and to S. (**C** and **D**) Visual discrimination performance of TcMAC21 and Eu measured by escape latency (p=0.27, C) and distance (p=0.19, D). Female: Eu (n=13), MAC21 (n=11); male: Eu (n=11), MAC21 (n=12). Data are analyzed by repeated measures ANOVA with LSD post-hoc and expressed as mean ± SEM. See also Data file S4.

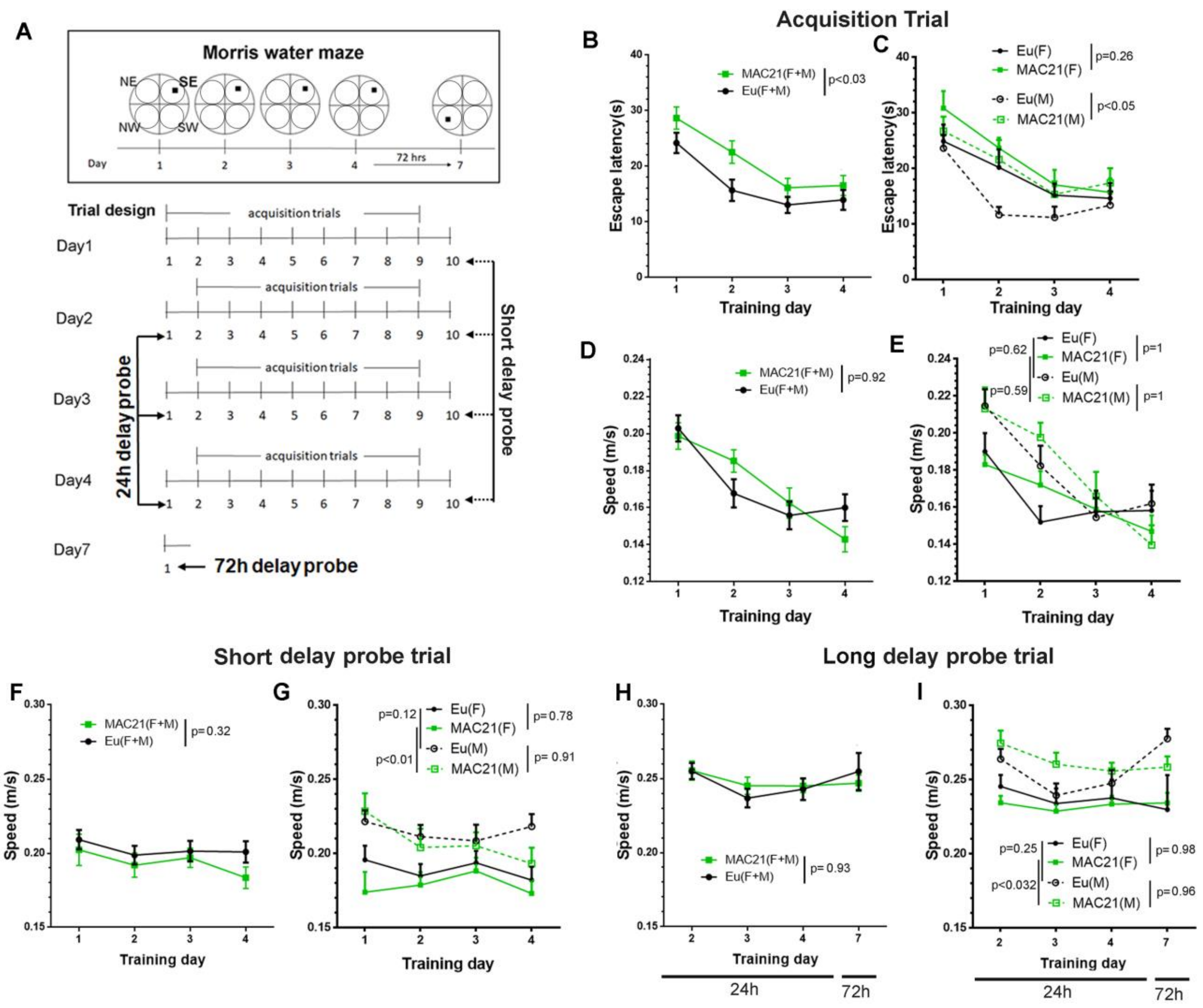


**Fig. S6. Classic MWM.** (**A**) Scheme for classic MWM: 4 full days of training, each day has 10 trials including 8 acquisition trials and 2 probe trials (one for ~30 min short delay and one for 24h long delay). A 72h delay probe trial is performed on day 7. (**B** and **C**) Escape latency in acquisition trials. (**D** and **E**) Swimming speed in acquisition trials. (**F** and **G**) Swimming speed in short delay probe trials. (**H** and **I**) Swimming speed in long delay probe trials. Female: Eu (n=10), MAC21 (n=11); male: Eu (n=11), MAC21 (n=12). Data are analyzed by repeated measures ANOVA with LSD post-hoc and expressed as mean ± SEM. See also Data file S4.

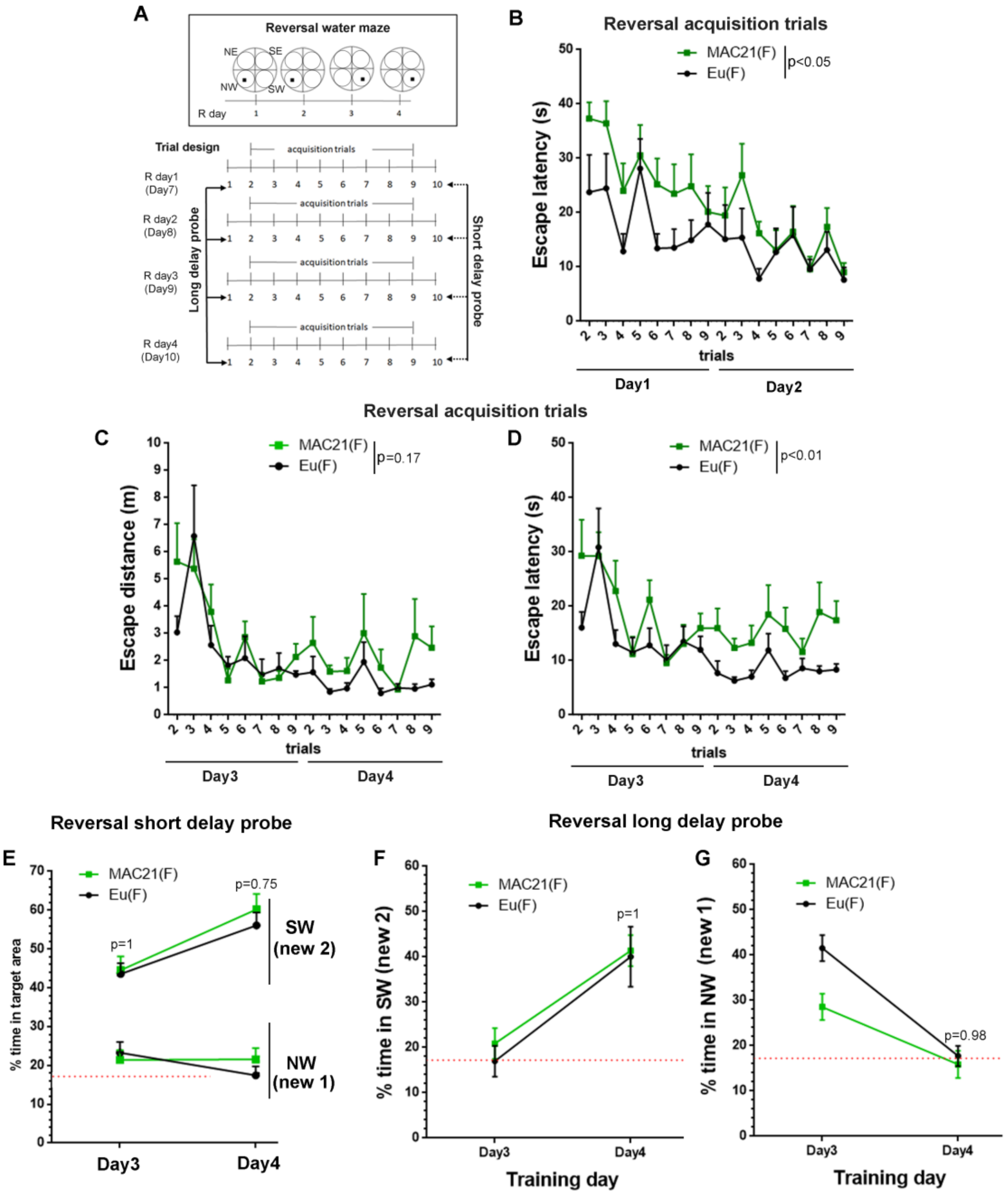


**Fig. S7. RRWM.** (**A**) RRWM scheme: 4 full days of training, the targeting platform is in NW during reversal 1 (R day 1 and R day 2) and relocated to SW during reversal 2 (R day 3 and R day 4). Trial 1 of R day 1 is known as the 72h delay probe trial of classic MWM. Each day has 10 trials, which includes 8 acquisition trials and 2 probe trials (one for ~30 min short delay and one for 24h long delay). (**B**) Escape latency in reversal 1. (**C** and **D**) Acquisition trials in reversal 2. (**E**) Short delay probe trials in reversal 2. (**F** and **G**) Long delay probe trials in reversal 2. Female: Eu (n=8), TcMAC21 (n=9)). Data are analyzed by repeated measures ANOVA with LSD post-hoc and expressed as mean ± SEM. See also Data file S4.

**Table S1. HSA21 genes were tested in TcMAC21 using human specific Taqman assay**

| human chromosome 21 Genes tested | Gene location | mouse chromosome | Taqman qRT-PCR |
|---|---|---|---|
| hTPTE (HSA21p) | 21p11.2 | | NO detection |
| hUSP25(deleted in MAC21) | 21q21.1 | 16 | NO detection |
| hJAM2 | 21q21.3 | 16 | Detected |
| hAPP | 21q21.3 | 16 | Detected |
| hRCAN1 | 21q22.12 | 16 | Detected |
| hDYRK1A | 21q22.13 | 16 | Detected |
| hBACE2 | 21q22.2 | 16 | Detected |
| hABCG1 | 21q22.3 | 17 | Detected |
| hHSF2BP | 21q22.3 | 17 | Detected |
| hC21orf2 | 21q22.3 | 10 | Detected |
| hPRMT2 | 21q22.3 | 10 | Detected |
| hS100B | 21q22.3 | 10 | Detected |

**Table S2. Absolute volume of brain structures in TcMAC21 and Eu were analyzed by MRI**

| Brain structures ($mm^3$) | WT (N = 7) | | MAC21(N=7) | | Mean difference (MAC21-WT) | Two-way ANOVA |
|---|---|---|---|---|---|---|
| | mean | std | mean | std | | P value |
| **Brain** | 467.27 | 14.62 | 487.86 | 17.55 | 20.59 | 0.0486 |
| **Hippocampus** | 23.69 | 2.14 | 24.69 | 1.20 | 1.00 | 0.1043 |
| **Cerebellum** | 63.45 | 3.76 | 62.29 | 4.97 | -1.16 | 0.3930 |
| **Lateral ventricles** (live imaging, n=3, t-Test) | 3.24 | 0.41 | 7.72 | 1.08 | 4.48 | 0.003 |
| Caudate putamen | 20.07 | 1.04 | 20.76 | 0.92 | 0.70 | 0.1888 |
| Neocortex | 91.82 | 8.35 | 94.01 | 8.99 | 2.19 | 0.5719 |
| External globus pallidus | 2.22 | 0.23 | 2.36 | 0.25 | 0.14 | 0.2114 |
| Superior colliculus | 8.78 | 0.63 | 10.15 | 0.50 | 1.37 | < 0.0001 |
| Thalamus | 19.56 | 1.26 | 20.97 | 1.20 | 1.41 | 0.0353 |
| Periform cortex | 3.21 | 0.19 | 3.21 | 0.24 | 0.00 | 0.8482 |
| Hypothalamus | 11.82 | 0.95 | 13.02 | 1.01 | 1.20 | 0.0134 |
| Amygdala | 6.75 | 0.42 | 7.11 | 0.33 | 0.36 | 0.0256 |
| Inferior colliculus | 3.38 | 0.26 | 3.66 | 0.22 | 0.28 | 0.0664 |
| Internal capsule | 4.55 | 0.51 | 4.79 | 0.28 | 0.24 | 0.2414 |
| Anterior commissure | 1.39 | 0.11 | 1.48 | 0.11 | 0.09 | 0.0705 |
| CC/External capsule | 13.53 | 0.92 | 13.98 | 1.25 | 0.45 | 0.3878 |
| Periaqueductal gray | 3.29 | 0.33 | 3.24 | 0.43 | -0.05 | 0.8527 |
| Septum | 1.15 | 0.51 | 1.18 | 0.62 | 0.03 | 0.7073 |
| Claustrum | 3.51 | 0.12 | 3.52 | 0.16 | 0.02 | 0.6270 |
| Nucleus accumbens | 2.03 | 0.17 | 2.23 | 0.20 | 0.20 | 0.0101 |

* Segmentation of brain, hippocampus, cerebellum and lateral ventricle were manually verified, and data were analyzed by two-way ANOVA.

**Table S3. Percentage of brain volume in TcMAC21 and Eu were analyzed by MRI**

| Brain structures | WT(N=7) mean(% of brain) | MAC21 (N=7) mean(% of brain) | Two-way ANOVA P value |
|---|---|---|---|
| **Hippocampus** | 5.07 | 5.06 | 0.8273 |
| **Cerebellum** | 13.58 | 12.76 | 0.0005 |
| **Lateral ventricles** (live imaging, n=3) | 0.66 | 1.52 | 0.005 |
| Caudate putamen | 4.29 | 4.26 | 0.8294 |
| Neocortex | 19.63 | 19.26 | 0.4333 |
| External globus pallidus | 0.47 | 0.48 | 0.6616 |
| Superior colliculus | 1.88 | 2.08 | **<0.0001** |
| Thalamus | 4.18 | 4.30 | 0.1281 |
| Periform cortex | 0.69 | 0.66 | 0.0021 |
| Hypothalamus | 2.53 | 2.67 | 0.0744 |
| Amygdala | 1.45 | 1.46 | 0.7722 |
| Inferior colliculus | 0.72 | 0.75 | 0.2883 |
| Internal capsule | 0.97 | 0.98 | 0.9869 |
| Anterior commissure | 0.30 | 0.30 | 0.4474 |
| CC/External capsule | 2.89 | 2.86 | 0.8851 |
| Periaqueductal gray | 0.71 | 0.67 | 0.3686 |
| Septum | 0.24 | 0.24 | 0.9624 |
| Claustrum | 0.75 | 0.72 | 0.1754 |
| Nucleus accumbens | 0.44 | 0.46 | 0.1653 |

* Segmentation of brain, hippocampus, cerebellum and lateral ventricle were manually verified, and data were analyzed by two-way ANOVA.

**Table S4. Peripheral blood analyses in TcMAC21 and Eu**

| Genotypes | (n) | RBC ($x10^4$/mL) | Hct (%) | Hb (g/dL) | MCV fL | MCH pg | MCHC % | Plt ($x10^4$/mL) | WBC ($x10^2$/mL) |
|---|---|---|---|---|---|---|---|---|---|
| WT | 6 | 1043 ± 47 | 49.2 ± 1.5 | 15.2 ± 0.5 | 47.2 ± 1.5 | 14.6 ± 0.5 | 31.0 ± 0.5 | 101.2 ± 6.8 | 181 ± 28 |
| MAC21 | 5 | 1009 ± 67 | 47.6 ± 3.3 | 14.9 ± 1.4 | 47.2 ± 0.8 | 14.8 ± 0.5 | 31.4 ± 0.8 | 112.3 ± 7.8 | 165 ± 14 |
| p-value | | $3.45 \times 10^{-1}$ | $3.14 \times 10^{-1}$ | $6.61 \times 10^{-1}$ | $9.92 \times 10^{-1}$ | $5.55 \times 10^{-1}$ | $3.45 \times 10^{-1}$ | $3.25 \times 10^{-2}$ | $2.88 \times 10^{-1}$ |

* Data were analyzed by two-tailed t-test and expressed as mean ± SD.

**Table S5. Husbandry in TcMAC21**

| Sire Genotype | Litters | Total | Litter size | MAC21 | MAC21 ratio | Male of total MAC21 pups |
|---|---|---|---|---|---|---|
| Euploid | 15 | 90 | 6 | 43 | 48% | 54% |
| MAC21 | infertile | - | - | - | - | - |

### Table S6. Primer sequences for genomic PCR or RT-PCR analyses

| | Gene or aim | Primer name (forward) | Forward primer(5'-3') | Primer name (reverse) | Reverse primer(5'-3') | Product size |
|---|---|---|---|---|---|---|
| | MAC1 | m11 5L | TGACAGAGAGCTTCCTCCTGCCTCTGTA | EGFP-F | CCTGAAGTTCATCTGCACCA | 5.0 kb |
| | Chr.21-loxP | #21CEN<1>2L | AAATGCATCACCATTCTCCCAGTTACCC | PGKr1 | GGAGATGAGGAAGAGGAGAACA | 4.5 kb |
| | Cre-loxP recombination | TRANSL1 | TGGAGGCCATAAACAAGAAGAC | TRANSR1 | CCCCTTGACCCAGAAATTCCA | 409 bp |
| | Cre-loxP recombination | kj neo | CATCGCCTTCTATCGCCTTCTTGACG | PGKr1 | GGAGATGAGGAAGAGGAGAACA | ~600 bp |
| | D21S265 | SHGC-40F | GGGTAAGAAGGTGCTTAATGCTC | SHGC-40R | TGAATATGGGTTCTGGATGTAGTG | 178 bp |
| | APP | SHGC-31514F | CTGGGCAATAGAGCAAGACC | SHGC-31514R | ACCCATATTATCTATGGACAATTGA | 115 bp |
| | D21S260 | D21S260F | AGCTGTTCATGCTTCCATCT | D21S260R | AGAGCCCAGAATATTGACCC | 270 bp |
| | SOD1 | SHGC-6902F | ATTCTGTGATCTCACTCTCAGG | SHGC-6902R | TCGCGACTAACAATCAAAGT | 133 bp |
| | D21S261 | SHGC-3610F | AACACCTTACCTAAAACAGCA | SHGC-3610R | TGGACCTTTTGATTTTTCCT | 130 bp |
| | AML1 | SHGC-30487F | GTAACCTGGTTAACATAGGGTTTC | SHGC-30487R | GTAGGGGAGGCTAATGGCAT | 150 bp |
| | CBR1 | J04056F | GATCCTCCTGAATGCCTG | J04056R | GTAAATGCCCTTTGGACC | 245 bp |
| | SIM2 | WI-22186F | GGGCCTCATGGTAAGAGTCA | WI-22186R | GAAAAATGTCGGTGGTATCTCC | 250 bp |
| | HLCS | WI-15188F | TTCAGTACCTCCCCAGATGC | WI-15188R | CTTAGTAGTGCAGACCTTTACCCC | 125 bp |
| | TTC3 | WI-19945F | TGGACAAATATAAGGCATGTTCA | WI-19945R | GTCACCTTCCTCTGCCTTTG | 267 bp |
| **Genomic PCR** | D21S394 | D21S394F | GGAGCCGGTTCTTCGAAGG | D21S394R | CAGCGTCCGGAATTCCTGC | 71 bp |
| | D21S336 | D21S336F | TCTGGTTCCCAGGATTGTAA | D21S336R | AGAGTTGCTGTAAGCATCAAAGT | 350 bp |
| | D21S55 | D21S55F | AGGCTCCTTCACCTCTTGAC | D21S55R | CATCCTCTTTGCATTAGG | 159 bp |
| | GIRK2 | GIRK2F | GTTTGTCTTCAGCTCACC | GIRK2R | CCCAAAATACTACACATCC | 266 bp |
| | ERG | M21535F | AATGGCGTCAGCCTCTCC | M21535R | CAGTTTGCCTTACGAGTGGTAGC | 254 bp |
| | ETS2 | SHGC-11267F | TACCATGCCAATGGTTTATAAGG | SHGC-11267R | ATGTGACTGGGAACATCTTGC | 177 bp |
| | D21S268 | D21S268F | CAACAGAGTGAGACAGGCTC | D21S268R | TTCCAGGAACCACTACACTG | 213 bp |
| | PCP4 | WI-14954F | GAATTCACTCATCGTAACTTCATTT | WI-14954R | CCTTGTAGGAAGGTATAGACAATGG | 126 bp |
| | D21S266 | D21S266F | CACAATGTAGATGTAGCACAGTTAG | D21S266R | TGAGTCTGAAGAAAGGCAAATGAAG | 166 bp |
| | D21S15 | D21S15F | GAGGATAAACCGATTCACAGCTAGGAATAC | D21S15R | GTGCACGTAATTAATGACCATGATATTGCT | 218 bp |
| | MX1 | WI-18875F | TGGACTGACGACTTGAGTGC | WI-18875R | CTCATGTGCATCTGAGGGTG | 143 bp |
| | TFF1 | SGC35308F | CAGGGATCTGCCTGCATC | SGC35308R | ATCGATCTCTTTTAATTTTTAGGCC | 183 bp |
| | PWP2 | SHGC-33273F | GATCTTGACCGGGAAAAGGG | SHGC-33273R | AACAAGTGGCAAAATGCATAC | 150 bp |
| | APECED | A009B16F | AAAATCCTCCCTTTAAGAGC | A009B16R | GGGTGTTAGGTACTGGCT | 118 bp |
| | PFKL | sts-X15573F | AGGGCTTCTGAGGCCAGC | sts-X15573R | AGGGCACTCTGTCCTCCTGC | 239 bp |
| | UBE2G2 | WI-11417F | TTCAACAGTCATTAGGTTCCACC | WI-11417R | GTGAGATCGGGAGAGGGAG | 129 bp |
| | APP | WI-18826F | ACGTTTGTTTCTTCGTGCCT | WI-18826R | GCCCCGTAAAAGTGCTTACA | 136 bp |
| | SOD1 | SHGC-6902F | ATTCTGTGATCTCACTCTCAGG | SHGC-6902R | TCGCGACTAACAATCAAAGT | 133 bp |
| | IFNAR2 | U29584F | CGAAGTTTCAGTCGGTGAG | U29584R | GGCATTCAGGTTTTATCCC | 181 bp |
| | TTC3 | WI-19945F | TGGACAAATATAAGGCATGTTCA | WI-19945R | GTCACCTTCCTCTGCCTTTG | 267 bp |
| **RT-PCR** | PCP4 | WI-14954F | GAATTCACTCATCGTAACTTCATTT | WI-14954R | CCTTGTAGGAAGGTATAGACAATGG | 126 bp |
| | MX1 | WI-18875F | TGGACTGACGACTTGAGTGC | WI-18875R | CTCATGTGCATCTGAGGGTG | 143 bp |
| | TFF3 | WI-7267F | GGCTGTGATTGCTGCCAG | WI-7267R | GTGGAGCATGGGACCTTTAT | 124 bp |
| | TFF1 | SGC35308F | CAGGGATCTGCCTGCATC | SGC35308R | ATCGATCTCTTTTAATTTTTAGGCC | 183 bp |
| | GADPH | RPC1 | CCATCTTCCAGGAGCGAGA | RPC2 | TGTCATACCAGGAAATGAGC | 722 bp |

**Dataset S1 (separate file).** WGS of TcMAC21

**Dataset S2 (separate file).** RNA-seq of HSA21 and mouse orthologs from TcMAC21 and Eu

**Dataset S3 (separate file).** Statistics tables of growth.

**Dataset S4 (separate file).** Statistics tables of behavioral tests
